# HESC-derived sensory neurons reveal an unexpected role for PIEZO2 in nociceptor mechanotransduction

**DOI:** 10.1101/741660

**Authors:** Katrin Schrenk-Siemens, Jörg Pohle, Charlotte Rostock, Muad Abd El Hay, Ruby M. Lam, Marcin Szczot, Shiying Lu, Alexander T. Chesler, Jan Siemens

## Abstract

Somatosensation, the detection and transduction of external and internal stimuli, has fascinated scientists for centuries. But still, some of the mechanisms how distinct stimuli are detected and transduced are not entirely understood. Over the past decade major progress has increased our understanding in areas such as mechanotransduction or sensory neuron classification. Additionally, the accessibility to human pluripotent stem cells and the possibility to generate and study human sensory neurons has enriched the somatosensory research field.

Based on our previous work, the generation of functional human mechanoreceptors, we describe here the generation of hESC-derived nociceptor-like cells. We show that by varying the differentiation strategy, we can produce different nociceptive subpopulations. One protocol in particular allowed the generation of a sensory neuron population, homogeneously expressing TRPV1, a prototypical marker for nociceptors. Accordingly, we find the cells to homogenously respond to capsaicin, to become sensitized upon inflammatory stimuli, and to respond to temperature stimulation.

Surprisingly, all of the generated subtypes show mechano-nociceptive characteristics and, quite unexpectedly, loss of mechanotransduction in the absence of PIEZO2.

## Introduction

Our ability to detect and transduce stimuli from the external and internal environment drives a major part of our daily behavior. The perception of innocuous mechanical stimuli for example, enables us to discriminate various textures and vibrations We can perceive a friendly touch on our arm as comforting and the warmth of the sun on our bare skin as pleasant. At the same time, we can perceive extreme temperatures, chemical compounds or mechanical forces as painful, leading to protective reflexive withdrawal responses to avoid tissue damage (Basbaum et al., 2009). The cells responsible for the detection and transduction of the respective stimuli are sensory neurons located in the dorsal root or trigeminal ganglia (DRG & TG). The plethora of stimuli that we are able to detect is reflected in the diversity of sensory neuron subtypes we have. Nociception, the ability to detect and transduce painful stimuli, is of special interest, as the primary function of pain as a warning system can outlive its usefulness by becoming a chronic state, which is a burden for the affected person. The pain-sensing neurons in the somatosensory system, the nociceptors, can, depending on their molecular make-up, respond to only one noxious stimulus such as temperature or to several stimuli such as noxious heat/chemicals as well as mechanical force (Basbaum et al., 2009). The latter are polymodal mechano-nociceptors.

Primary human sensory neurons are difficult to access and therefore did not have a major role as model system to study human nociception. This has changed tremendously with the accessibility of human embryonic and induced pluripotent stem cells (hESCs & hiPSCs), which now offers the unprecedented opportunity to derive sensory neurons from pain afflicted patients. Certainly, this depends on suitable, efficient procedures to differentiate the cells. A few protocols have been published, describing how to generate sensory neurons pursuing different strategies: either small molecule inhibitors (Chambers et al., 2012) are used to drive stem cells towards sensory neuron fate or virally induced overexpression of two to five transcription factors directly reprogram human fibroblasts or stem cells to become sensory neurons (Blanchard et al., 2015; Wainger et al., 2015).

We have chosen a more classical approach, by generating the progenitor cells of sensory neurons –neural crest cells– as starting material for subsequent differentiation. Exploiting this strategy, we have already been able to generate functional mechanoreceptors and successfully showed their usefulness as a tool to study mechanotransduction *in vitro* (Schrenk-Siemens et al., 2015). In the present study, we again used hESC-derived neural crest-like cells as a starting population for our differentiation procedure and virally induced the expression of *NEUROGENIN1* (*NGN1*). NGN1 is a basic helix loop helix factor critical for nociceptor development in rodents (Ma et al., 1999). The differentiation process was further induced by the addition of morphogens and neurotrophic factors.

Native nociceptors are not a homogenous cell population, rather, they are molecularly diverse, reflecting their ability to respond to different types of stimuli and differentially mediate and modulate painful responses (Pace et al., 2018). Within our differentiation paradigm we diversified the generated nociceptors by varying the duration of *NGN1* expression and by exposing the differentiating cells to different extracellular factors. Using different genetically engineered hES cell lines and analytical techniques, such as RNAseq, cytochemistry, functional calcium-imaging and electrophysiology, we characterized the resulting cell populations and compared them to cultured native rodent sensory neurons and previously reported human-derived nociceptors.

The thereby generated neurons showed molecular and functional characteristics of nociceptors. One differentiation protocol in particular generated a homogenous population of sensory neurons expressing TRPV1, a prototypical molecular marker of nociceptors and a major driver of inflammatory pain (Caterina et al., 2000). Accordingly, we find the cells to homogenously respond to capsaicin and to have the potential for becoming sensitized upon inflammatory stimulation Both of these characteristics are hallmarks of native nociceptors and a notorious feature that has been discussed to drive/fuel pathological forms of pain (Pace et al., 2018; Reichling and Levine, 2009). Additionally, these neurons respond to temperature stimulation, in agreement with the expression of multiple thermosensitive TRP ion channels, including TRPV1.

Similar to the abundance of mechanosensitive nociceptors in DRG cultures of the mouse, many of the derived cells were inherently mechanosensitive. Furthermore, many of the differentiated neurons expressed PIEZO2, a protein recently identified as key driver of mechanotransduction in low threshold mechanoreceptors in mice and humans (Chesler et al., 2016; Ranade et al., 2014; Schrenk-Siemens et al., 2015). To our surprise, deletion of the *PIEZO2* gene abolished mechanotransduction in the hESC-derived nociceptors. Analogue experiments using cultured mouse nociceptors also abolished mechanically evoked currents in the absence of PIEZO2, suggesting that this receptor is also involved in mechanotransduction in a selective subset of mouse and human nociceptive neurons *in vitro*.

In summary, we here describe the comparison of several novel procedures for the generation of diverse populations of human nociceptor-like cells, with the emphasis on one particular protocol, which enables the efficient production of a homogenous population of TRPV1-expressing neurons, showing molecular and functional characteristic hallmarks of pain-sensing neurons. To our knowledge, none of the previously published protocols describes a similar nociceptor population, that has been characterized and compared in such depth and that can serve as a translational tool to study humane (mechano-) nociception *in vitro*.

## Results

### Generation of hESC-derived *TRKA*-positive neurons

The richness of sensations the human body can experience is the result of a plethora of different sensory neurons, which are specialized to detect and transduce one or several distinct stimuli. The diverse spectrum of sensory neurons emerges from a transient cell population during development: the neural crest cells, which serve as progenitors for the sensory neurons located in the dorsal root ganglia (DRG). How the multitude of different sensory subtypes is generated is still not fully understood, however, a combination of extracellular factors and intrinsic cell-autonomous programs, such as the orchestrated expression of different transcription factors, are major driving forces (Kalcheim and Kumar, 2017). In particular, early instructive roles of Neurogenin1 (NGN1) and Neurogenin2 (NGN2), two transcription factors expressed in sensory lineage-primed neural crest cells, are well-documented (Ma et al., 1999). While the majority of large diameter TRKB-and TRKC-positive sensory neurons primarily depend on NGN2, the generation of small diameter neurons requires NGN1 expression. It is this latter group that comprises nociceptors, sensory neurons specialized for detecting painful – often considered harmful– stimuli.

We made use of this defining feature and transiently expressed *NGN1* in hESC-derived neural crest-like cells (NCLCs), a cell population that we and others have previously shown to inherently hold the potential to become sensory-like neurons (Bajpai et al., 2010; Schrenk-Siemens et al., 2015). In order to drive and control *NGN1* expression, NCLCs were infected with two lentiviruses (Fig 1A; Fig EV1A): one lentivirus expressed a tetracycline-controlled transactivator (rtTA) under the control of an ubiquitin promoter for constitutive expression of rtTA. The second lentivirus expressed *NGN1*, *EGFP*, and a puromycin resistance cassette linked with P2A and T2A sequences, respectively– under the control of the tetracycline operator (tetO) promoter. By adding doxycycline (Dox) to the culture medium, the expression of *NGN1* was induced and verified by the presence of EGFP. Thereby we were able to reversibly control the duration of *NGN1* expression (Fig 1A, Fig EV1B). Interestingly, after switching viral *NGN1* expression off, *NGN1* transcription initially declined, but stayed elevated in comparison to baseline levels. Notably, *NGN2* was not detectable at any time point (Fig EV1C). Presumably, the endogenous *NGN1* gene (but not the *NGN2* gene) was activated in differentiating cells.

**Figure 1.**
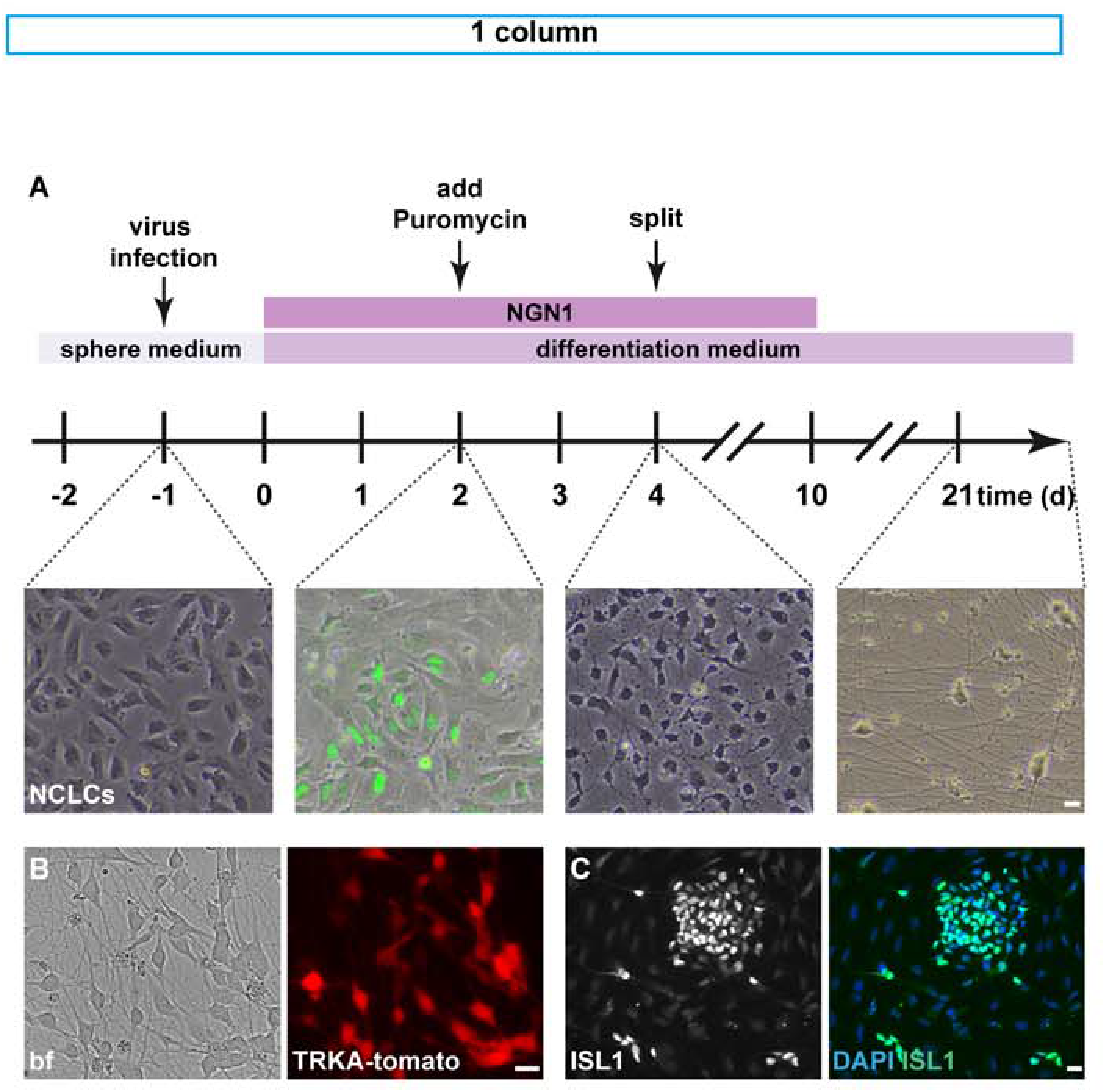
Generation of hESC-derived nociceptor-like cells. A. Schematic illustration of the differentiation protocol to generate hESC-derived nociceptor-like cells: arrows indicate time point of virus infection, selection (addition of puromycin) and splitting of cells. Graphs show characteristic morphology of cells at the indicated time points: −1 day (d) neural crest-like cells (NCLCs), 2 d *NGN1* expression, indicated by GFP-signal; 4d neural progenitors show small neurites; ≥ 21d neurons with long processes can be observed. B. Live cell imaging of neuronal progenitors derived from the hESC TRKA-tomato reporter cell line: left picture shows a bright field (bf) image, right picture the tomato signal of cells 6 days after doxycycline-induced expression of *NGN1*. C. Immunostaining of hESC-derived NCLCs 2 days after induction of *NGN1*-expression. As early as 2 days after *NGN1* induction the presence of the pan-sensory marker ISLET1 (ISL1) is detectable in a subset of cells. (DAPI: blue; ISL1: green). Scale bars: 20μm.

One to two days after onset of exogenous *NGN1* induction, the NCLCs began to change their morphology and islands of roundish EGFP-positive cells, encircled by elongated, EGFP-negative cells could be observed (Fig 1A). To eliminate non-infected (EGFP-negative) cells and to enrich *NGN1* expressing NCLCs, puromycin was added for up to 48 hours into the Dox containing culture medium. Four to five days after induction of *NGN1* expression, neuronal progenitors with small neurites were observed (Fig 1A), most –if not all– expressed *NGN1*/EGFP (data not shown).

*In vivo, NGN1* is transiently expressed in neurons that will express the receptor tyrosine kinase *TRKA*, a gene that marks nociceptive sensory neurons (Ernsberger, 2009). We therefore limited expression of *NGN1* to the first 10 days of differentiation (Fig 1A).

Morphological features alone are insufficient to unambiguously demarcate nociceptive neurons in mixed neuronal cultures. Therefore, we generated a hESC reporter line, which expresses td-tomato under the control of gene-regulatory elements of the TRKA receptor. The *TRKA* gene was left intact and the tomato sequence was introduced before the Stop codon of the *TRKA* locus with a T2A linker sequence using CRISPR/Cas9 technology (Fig EV2A–B). The fluorescent reporter allowed us to follow the *in vitro* differentiation of nociceptive neurons by microscopy: Six days after *NGN1* induction, the first tomato-positive cells could be observed when using the TRKA reporter cell line (Fig 1B). The tomato positive neurons started to develop processes shortly after *NGN1* induction and continued to generate an elaborate network of neuronal processes in the course of the culturing period (Fig 1A).

Two days after *NGN1* induction, the NCLCs started to express ISLET1 (Fig 1C), an indication that the cells had begun to acquire sensory neuron-like fate. Puromycin selection yielded sensory neuron cultures of high purity (84.6 ± 12,6 % after 21-25 days *in vitro* (div); 90.9 ± 4,6% after 42-58 div). Cultures not virally induced to express *NGN1* did not express the fluorescent TRKA reporter (data not shown).

To further assess the validity of the TRKA reporter and to verify co-expression of the fluorescent tomato transgene together with endogenous *TRKA* transcripts, double fluorescence *in situ* hybridizations were performed. These experiments demonstrated high (close to 100%) overlap of tomato-and *TRKA* expression (Fig EV2C).

Collectively, these results show that driving expression of *NGN1* in hESC-derived sensory neuron precursors (NCLCs) is sufficient and highly efficient in generating *TRKA*-positive **<u>no</u>**ci**<u>c</u>**eptor-**l**ike neurons that from here on we dub NOCL neurons.

### Functional characterization of *TRKA*-positive neurons

Mature nociceptors are specialized to detect a variety of disparate physical (mechanical and thermal) as well as inflammatory stimuli, which accounts for differences in their molecular armament (Li et al., 2016; Usoskin et al., 2015). These differences likely are the result of developmentally-induced differences in their transcriptional programs (Lallemend and Ernfors, 2012). Previous attempts to generate nociceptors have shown that it is difficult to efficiently produce discrete nociceptor subtypes (Blanchard et al., 2015; Chambers et al., 2012; Wainger et al., 2015). We therefore wondered whether recapitulating key steps of nociceptor development –instead of trans-differentiating hESCs or fibroblasts directly into nociceptors as employed in previous protocols– would create a permissive environment to yield homogenous populations of discrete nociceptor subtypes. With this goal in mind, we first generated neural crest-like cells (NCLCs), altered their internal transcriptional program and varied the composition of external (neurotrophic) factors.

During mouse DRG development NGN1 is transiently expressed. However, it is unclear at what time point during human DRG development *NGN1* is turned on and for how long its expression persists. We therefore compared conditions of transient and extended *NGN1* expression designated NOCL1 and NOCL2, respectively (Fig 2A). Additionally, external factors such as neurotrophins have been shown to influence sensory neuron fate. We therefore included a culturing condition devoid of serum, instead, we used a defined medium only containing neurotrophins implicated in sensory neuron differentiation (NOCL3-Fig 1A and 2A; see methods for details). Last but not least, we also modified a previously published differentiation protocol and included two additional transcription factors *KLF7* and *ISL2* –instead of *NGN1* alone– which had been shown to induce nociceptor-like cells (Wainger et al., 2015). Different to the previous protocol, which uses human fibroblasts or embryonic stem cells as starting cell population, we introduced these factors into derived NCLCs, based on the assumption that the native precursors of nociceptors are more readily primed to differentiate into sensory neurons. We refer to this differentiation procedure as NOCL4 (Fig 2A).

**Figure 2.**
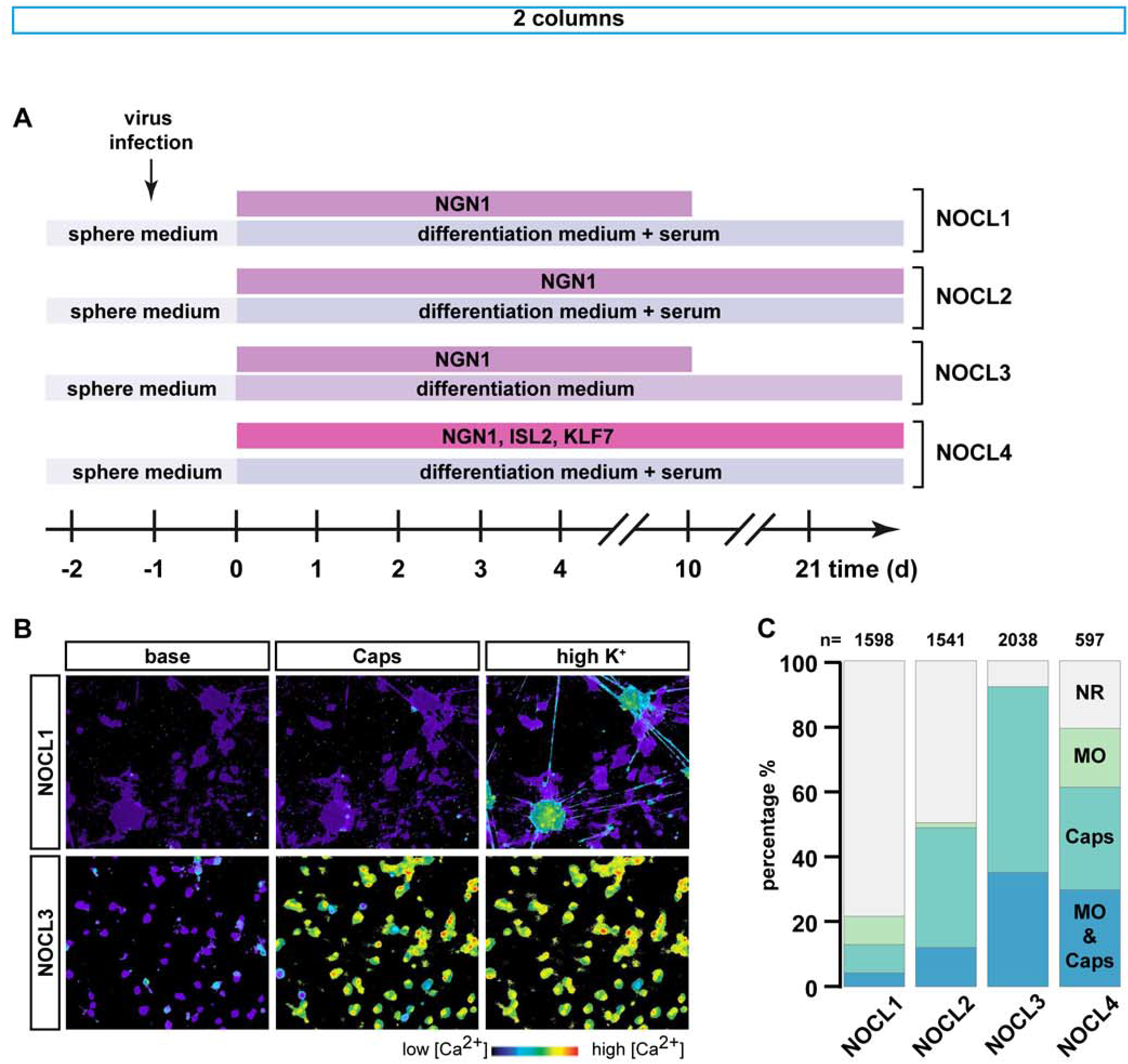
Functional characterization of different NOCL-types. A. Schematic illustration of the differentiation procedures to generate different types of hESC-derived nociceptor-like cells (NOCL1–4), showing the differences in medium composition (± serum), the type of transcription factors used (*NGN1*, *ISL2* and *KLF7*) and the duration of their expression (10 days or until the end of the culturing period). B. Representative pseudo-colored images of calcium responses observed in NOCL1 and NOCL3 cells loaded with Fura2 before (base) and during either stimulation with capsaicin (Caps, 1μM) or high K^+^ Ringer solution. C. Stacked bar charts, showing the percentage of NOCL1–4 cells responding to mustard oil (MO, 200μM), capsaicin (1μM), or both agonists using calcium imaging. For NOCL1: 78.4 ± 3% non responders, 8.7 ± 2.5% MO responders, 8.7 ± 1.9% Caps responders, 4.2 ± 1.6% MO and Caps responders (N=6). For NOCL2: 49.9 ± 7.3% non responders, 1.6 ± 0.3% MO responders, 36.9 ± 9.6% Caps responders, 12.1 ± 4.3% MO and Caps responders (N=4). For NOCL4: 20.7 ± 6.7% non responders, 18.0 ± 11.3% MO responders, 31.5 ± 11.1% Caps responders, 29.7 ± 8.2% MO and Caps responders (N=3). For NOCL3: 8.0 ± 5.9% non responders, 0% MO responders, 57.1 ± 15.0% Caps responders, 35.1 ± 16.6% MO and Caps responders (N=4). Data shown as average percent ± SEM.

Next to *TRKA*-expression, a hallmark of many nociceptors is the expression of the capsaicin receptor TRPV1 (Basbaum et al., 2009). Thus, a common way to identify and characterize nociceptors is to test their response to capsaicin, the pungent ingredient of chili peppers that triggers the opening of the TRPV1 ion channel to permit cation permeation such as the influx of calcium ions. Accordingly, following a differentiation period of at least 3 weeks, we evaluated the differently generated NOCLs for their ability to respond to capsaicin. NOCL cultures were loaded with the calcium indicator Fura-2 and challenged with 1μM capsaicin. Remarkably, we found that the NOCL cultures differed considerably in their response to Capsaicin: While only 13 ± 2% of neurons responded to capsaicin when *NGN1* was turned off after 10 days (NOCL1 differentiation protocol; Fig 2B–C), expressing *NGN1* constitutively for the entire differentiation period resulted in 48,9 ± 7,6% of capsaicin responders (Fig 2C). Cultures derived using a cocktail of virally expressed transcription factors *NGN1*, *ISL2*, and *KLF7* (Wainger et al., 2015) (NOCL4) produced 61,1 +/-14,7% capsaicin responsive neurons (Fig 2C). Strikingly, limiting *NGN1* expression to 10 days and nurturing differentiating neurons in a defined culture medium absent of any serum (NOCL3 - see methods for details) resulted in a nociceptor-like cell population that homogenously responded to capsaicin: We found that most if not all derived neurons responded to capsaicin (92,2 ± 5,9%). This is in stark contrast to low threshold mechanoreceptors (LTMRs), generated using our previously published protocol (Schrenk-Siemens et al., 2015), which did not yield any TRPV1-expressing neurons, and that here served as a control cell population (Fig 2B–C and Fig EV3A).

The ability to efficiently generate TRPV1/*TRKA*-positive NOCL neurons was specific for *NGN1* and could not be recapitulated by viral expression of either *RUNX1* (Fig EV3B) or *TRKA* (Fig EV3C), two genes that are induced subsequent to NGN1 expression during murine nociceptor development (Lallemend and Ernfors, 2012).

In native mouse and human DRGs, another TRP ion channel, TRPA1, is expressed in a subset of TRPV1-positive neurons (Rostock et al., 2018). Interestingly, the differentiation protocols also differed considerably in the fractions of TRPA1-postive neurons as assessed by calcium imaging using the TRPA1 agonist mustard oil: NOCL1 neurons showed only a small population of TRPV1 cells responding to mustard oil (4,2 ± 1,6%) (Fig 2C). In NOCL2-, NOCL3- and NOCL4 cultures, fractions of 12,1 ± 4,3 %, 35,1 ± 16.6% and 29,7 ± 8,2% of capsaicin-responsive cells responded to mustard oil, respectively (Fig 2C). Almost none of the cultures harbored any menthol responsive cells (data not shown). Collectively, these results show that the four NOCL differentiation procedures yielded different fractions of sensory TRP channel positive neuronal populations.

### Molecular characterization of *TRKA*-positive NOCL neurons

To gain a more complete picture of the molecular identity of the derived nociceptor cells, we surveyed gene expression at several time points throughout the differentiation period (day 0 –before *NGN1*-induction–, day 12, day 24 and day 31), focusing on the NOCL3 protocol yielding the highest percentage of *TRKA*/TRPV1-positive neurons (Fig 2C). We found that *NGN1* expression triggered a gene program characteristic of developing sensory neurons, with *ISLET1* and *BRN3A* induced early on during the differentiation process (Fig 1C and 3A, Fig EV4). Expression profiling showed that, next to TRPV1, other prototypical nociceptor genes such as NA_V_1.7, NA_V_1.8 and NA_V_1.9 (Habib et al., 2015) were expressed upon NOCL3 differentiation (Fig 3A–B, Fig EV4). We also found strong induction of *TRKA* expression (Fig 3A, Fig EV4), further confirming the reliability of our TRKA-tomato reporter cell line. Transcript-levels for most of these nociceptor markers reached a stable expression level between 24 to 30 days of differentiation (Fig 3A–B and Fig EV4), suggesting that a steady state was reached.

**Figure 3.**
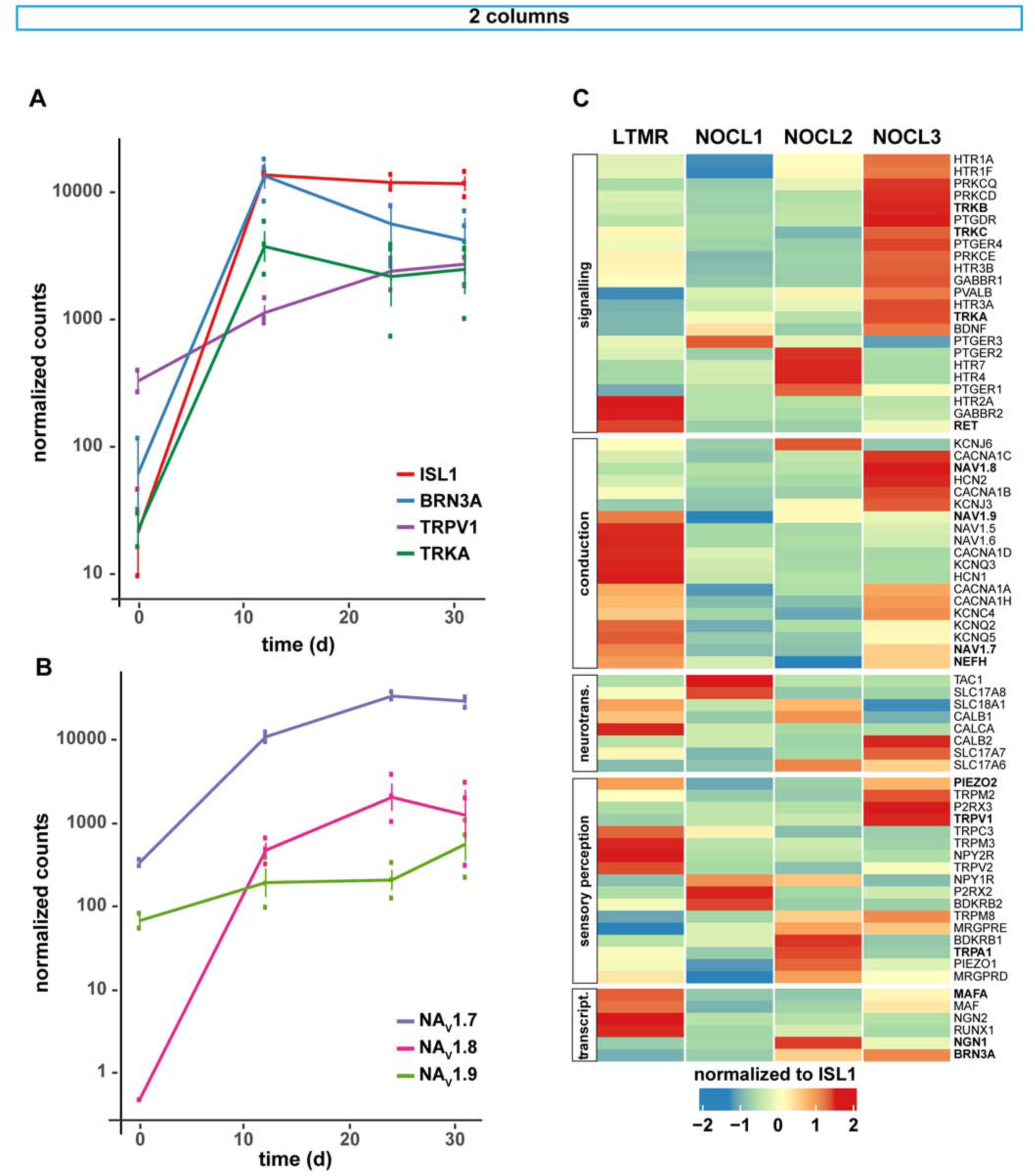
Transcriptome analysis of different hESC-derived sensory neuron subtypes. A. RNA-seq analysis of *ISL1*, *BRN3A*, *TRPV1* and *TRKA* transcripts in NOCL3 cells at specific time points during their differentiation (0, 12, 24 and 31 d), shown as normalized counts (for analysing details see material and methods; N=3). B. RNA-seq analysis of NA_V_1.7, NA_V_1.8 and NA_V_1.9 transcripts in NOCL3 cells at specific time points (0, 12, 24 and 31 d) during their differentiation, shown as normalized counts (N=3). C. Heatmap showing the relative expression of selected genes in LTMR and NOCL1–3 cells after 24 days of differentiation. Genes are categorized into functionally connected groups and those indicated in bold have been analysed in more detail (neurotrans. = neurotransmission, transcript. = transcription). Gene counts were first normalized and then related to the pan-sensory marker ISL1 and averaged (N=3). Depicted are center-scaled values.

We wondered whether subtly altering the *in vitro* differentiation procedure (duration of *NGN1* induction, availability of external morphogenetic factors) would change the fate of the ensuing NOCL neurons. Therefore, we compared expression profiles of the different NOCL cultures at 24 days in culture, a time point, where a steady state for several key nociceptive genes was observed (Fig 3A–B).

Indeed, we found that prolonging the expression of *NGN1* for the entire duration of the differentiation period (NOCL2) or enriching the medium with serum factors (NOCL1 and NOCL2) had a strong impact on the emerging sensory neurons: short (but not long) *NGN1* expression induced the appearance of TAC1 (Substance P), a marker of peptidergic nociceptors, in NOCL1 neurons (Fig 3C). TRPV1, the capsaicin receptor, present in both, peptidergic and non-peptidergic nociceptors (Dirajlal et al., 2003; Li et al., 2016; Usoskin et al., 2015), was robustly expressed in *TRKA*-positive NOCL3s while largely absent from LTMRs and was expressed significantly lower in serum-derived NOCL1s and NOCL2s (Fig 3C), matching the unequal capsaicin responses observed in the different NOCL1-3 cell types (Fig 2B–C).

Other genes of functional relevance for nociceptors and implicated in their sensitization, including G-Protein-coupled receptors (e.g. Serotonin-, Prostaglandin-and Histamin receptors), essential signaling molecules (such as *PKCε/PKCδ*), and *P2X2/3*- and *HCN* ion channels were also found to be differentially expressed in NOCL1-3 cultures (Fig 3C).

We found the 3 TRK receptors *TRKA*, *TRKB* and –to a lesser extent– *TRKC*, co-expressed in the NOCL3 neuron population while NOCL1- and 2 neurons were largely devoid of *TRKB/C* (Fig 3C and Fig EV2C and EV5A–C). In native murine DRG neurons, conjoint TRK expression is observed during DRG neuron development (Ernsberger, 2009), while in mature sensory neurons, TRKB and TRKC have classically been associated with LTMRs, proprioceptors and other non-nociceptive sensory subtypes. Somewhat surprisingly, we recently reported *TRKB* and *TRKC* to be expressed in a small fraction of native *TRKA*- positive small-diameter neurons in human DRGs (5 and 16%, respectively) (Rostock et al., 2018). It is therefore possible that the NOCL3 population we describe here either represents a discrete nociceptor subtype that expresses multiple TRK receptors or, alternatively, these cells have not yet reached full maturity.

While the level of several transcripts expressed in mature nociceptors plateaued towards the end of the differentiation period (Fig 3A–B), NA_V_1.9, an ion channel associated with pathological forms of pain, had not reached a steady state at the end of the differentiation period but started to increase expression in the 2^nd^ half of the differentiation phase (Fig 3B, Fig EV4), providing evidence for the latter possibility and suggesting that NOCL3 neurons are not yet fully mature.

It is unclear to what degree insights gleaned from murine models can be extrapolated to human nociceptors. While the overall molecular profile of human and rodent nociceptors is similar, significant molecular differences of *discrete* nociceptor subtypes have been observed (Rostock et al., 2018). One prominent difference between native human vs. murine nociceptors is the presence of neurofilament heavy (NEFH, NF200) in most –if not all– human DRG neurons (Rostock et al., 2018), a protein selectively expressed in myelinated murine Aδ- and β-fibers (Basbaum et al., 2009). In agreement with these findings, we find NF200 to be expressed in all NOCL neurons, regardless of the differentiation protocol used (Fig 4A).

**Figure 4.**
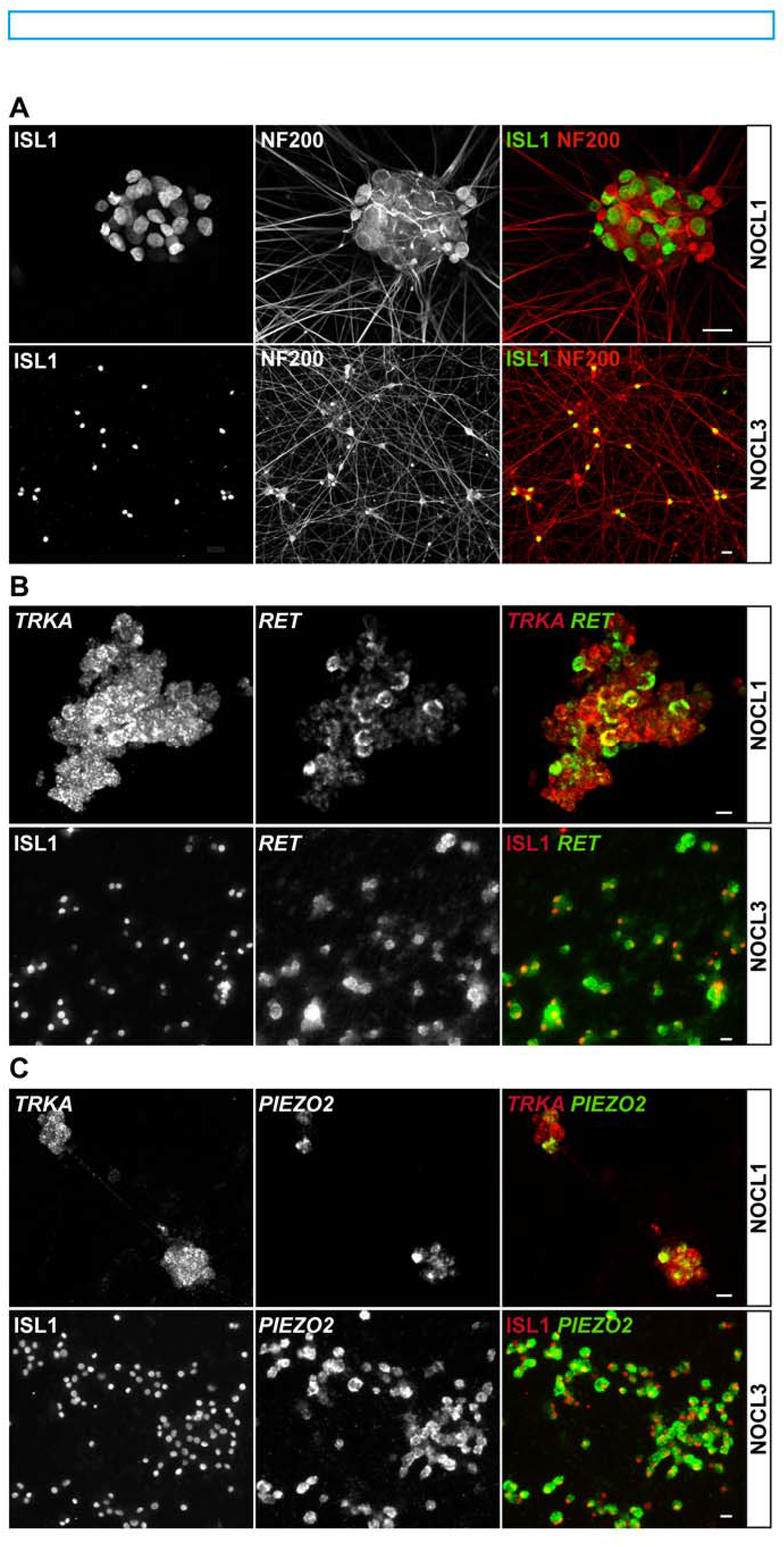
Molecular characterization of NOCL1 and NOCL3 neurons. A. Immunostaining of NOCL1 and NOCL3 cells detecting ISL1 protein (left), NF200 (middle), and the composite image (right). Note, the presence of neurofilament heavy 200 is essentially detected in all NOCL cells. B. *In situ* hybridization of NOCL1 and NOCL3 cells with probes detecting *TRKA* (upper left and red) and *RET* (middle and green) as well as immunostaining for ISL1 (lower left and red) to show co-expression of *TRKA* and *RET* or ISL1 and *RET*. In NOCL1 cells 77.6 ± 0.5% of *TRKA* positive neurons co-expressed *RET*, compared to 97.4 ± 0.9% in NOCL3 cells (N=3). C. *In situ* hybridization of NOCL1 and 3 cells with probes detecting *TRKA* (upper left and red) and *PIEZO2* (middle and green) as well as immunostaining for ISL1 (lower left and red) to show co-expression of the respective markers: 64.2 ± 2.3% of *TRKA* positive NOCL1 neurons co-expressed *PIEZO2*, compared to 95.0 ± 2.4% in NOCL3 cells (N=3; mean ± SEM; scale bar 20μm).

The receptor tyrosine kinase RET is robustly expressed in both murine and human DRGs and associated with different classes of sensory neurons. In rodents RET is expressed in a graded fashion in different subsets of nociceptors, itch-detecting neurons and LTMRs (Stantcheva et al., 2016) but largely excluded from TRKA-positive neurons (Franck et al., 2011). However, in human neurons, RET is differently distributed with nearly double the fraction of mature human nociceptors co-expressing *TRKA* and *RET* (mouse: 23,2 +/- 1,8 % vs. human: 45,9 +/- 0,7 %, Rostock et al., 2018). We found *RET* to be robustly expressed in *TRKA*-positive NOCL3 neurons (97,4 ± 0,91%), while the receptor was found in 77,6 ± 0,5% in *TRKA*-positive NOCL1 neurons (Fig 4B). The expression pattern clearly coincides more closely to what has been shown in native human DRG neurons –but only rarely in mouse DRG neurons–*TRKA* expression does not preclude the expression of *RET* in mature human nociceptors (Rostock et al., 2018; Stantcheva et al., 2016).

While RET is widely expressed across small- and large diameter neurons, the transcription factor MAFA is a more selective molecular marker for mouse and human LTMRs and largely excluded from nociceptors (Wende et al., 2012). We find *MAFA* transcripts absent from NOCL1 and NOCL2 neurons but detectable in NOCL3 neurons, albeit to a lesser extent compared to LTMRs (Fig EV5D). In agreement with a minor –if any– function for MAFA in nociceptors, we find that MAFA protein is only detectable in a few NOCL3 neurons, differing from derived LTMRs which robustly express MAFA (Fig EV5E).

Another sensory receptor, PIEZO2, previously identified as the main transducer of mechanical stimuli in LTMRs (Ranade et al., 2014; Schrenk-Siemens et al., 2015), was expressed in NOCL neurons, in agreement with data obtained from native mouse and human DRG tissue (Rostock et al., 2018). All NOCL differentiation procedures yielded *PIEZO2*-postive neurons, albeit the expression level and the fraction of neurons positive for the ion channel varied (Fig 3C and Fig 4C).

In summary, driving *NGN1* expression in human neural crest-like cells combined with exposing them to different extracellular factors, promoted the generation of different nociceptor-like cells, with similar molecular profiles compared to subclasses of native DRG neurons. We found that NOCL1 cells were more reminiscent of peptidergic nociceptors while NOCL3 cells expressed several genes typically found in non-peptidergic nociceptors (Li et al., 2016; Usoskin et al., 2015) (Fig EV6). NOCL protocols were also successful in converting human iPS cells into nociceptors (data not shown).

### NOCL3 neurons are heat responsive and can be sensitized by inflammatory stimuli

A number of TRP ion channels have been implicated in mediating responses from warm to hot temperatures (Siemens and Kamm, 2018). In DRG sensory neurons, the most prominent warmth/heat sensitive TRPs are TRPV1, TRPM2, TRPM3, and TRPA1 (Tan and McNaughton, 2016; Vandewauw et al., 2018; Yarmolinsky et al., 2016). Therefore, we next tested the response of the generated sensory neurons to temperature steps from 35°C to 43°C using Ca^2+^-imaging focusing on the population of NOCL3 cells that featured the highest percentage of TRPV1 responders. Different to cultured mouse DRG neurons, hESC-derived neurons responded more homogenously to the temperature stimuli (Fig 5A). These results corroborated uniformity of cells produced by the NOCL3 protocol, resulting in a population of prototypical TRPV1-positive, temperature responsive nociceptors.

**Figure 5.**
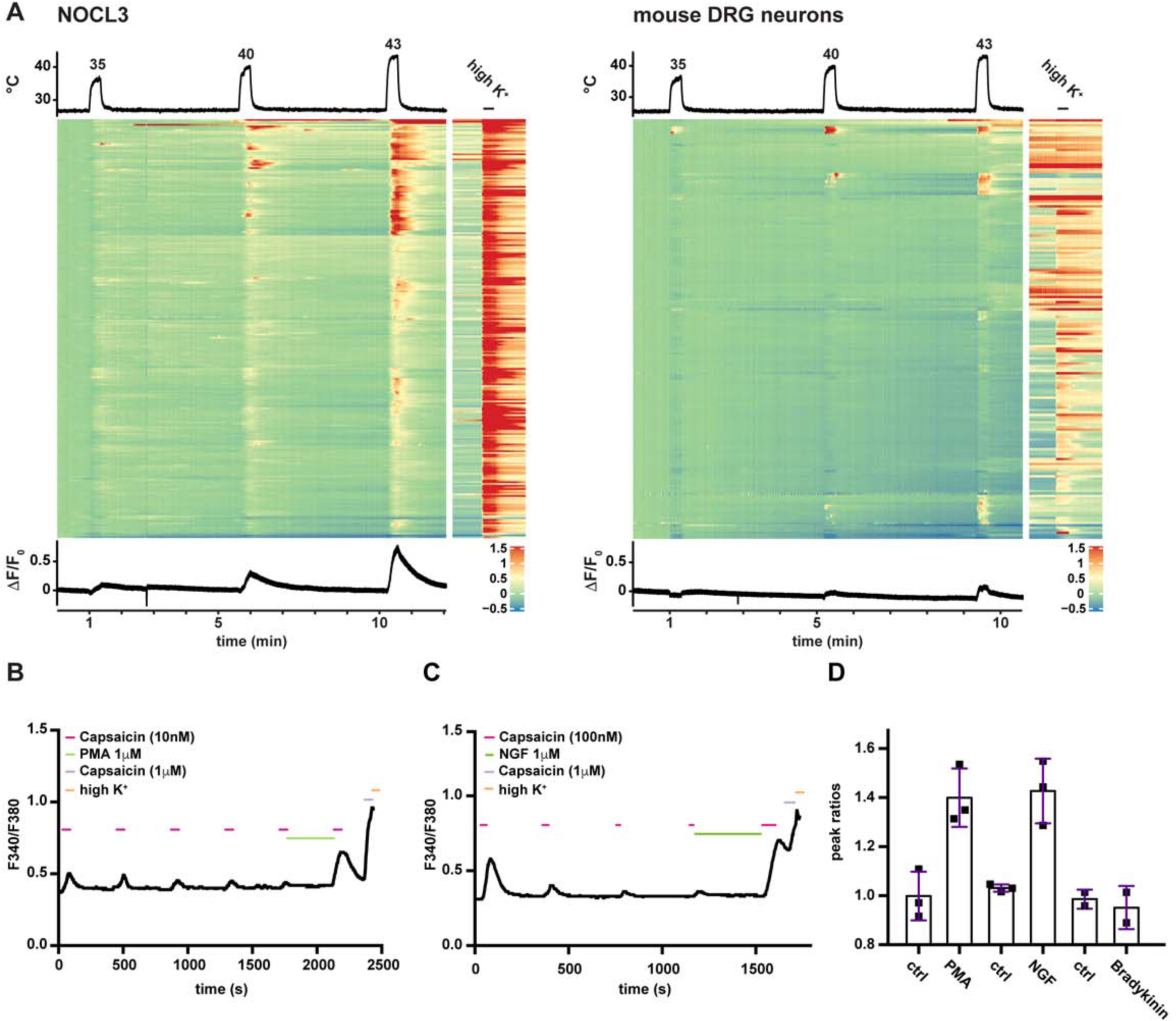
Nociceptor sensitization assessed by calcium-imaging. A. Heatmaps of calcium imaging experiments challenging Cal520 loaded NOCL3 and mouse dorsal root ganglion (DRG) neurons with increasing temperatures from 35°C to 43°C and subsequently with high K^+^ Ringer solution (to identify all neurons). Every row represents a single cell. Depicted is the ΔF/F_0_ ratio using the baseline before the first stimulus as F_0_. The trace below the heat map shows the average response of all neurons (NOCL3 n=248 cells; DRG n=174). B. Representative fluorescence ratios (340nm/380nm) obtained from one NOCL3 neuron loaded with Fura2 and stimulated with multiple pulses of capsaicin (10nM). Incubation of cell with the phorbol ester PMA (1uM) for 5 min results in sensitization leading to an increased response amplitude in response to a low concentration of capsaicin. Subsequent application of high K^+^ ringer solution showed presence of all functional neurons. C. Calcium response of a NOCL3 neuron stimulated repetitively with capsaicin (100nM). Incubation with NGF (100ng/ml) for 5 min sensitized the cell, resulting in an increased response to capsaicin. High K^+^ ringer solution was added at the end to visualize functionality of the neuron. D. Bar graphs represent the ratios between the peak response after- and the peak response before application of inflammatory agents as indicated. Fraction of cells showing sensitization: PMA: 91% (753 of 823 cells); NGF: 28% (173 of 611 cells); bradykinin: 0% (0 of 319 cells).

Besides responding to heat, the capsaicin receptor can also integrate inflammatory stimuli. Inflammatory sensitization is a characteristic feature of TRPV1. We therefore investigated NOCL3 neurons in the context of inflammatory pain signaling.

Components of the so called ‘inflammatory soup’, a mixture of signalling molecules such as Bradykinin, NGF, Serotonin and others are released at the site of inflammation and activate disparate signalling cascades present in different subsets of nociceptors. These inflammatory cascades converge on TRPV1 and boost channel activity to subthreshold stimuli (primary hyperalgesia) (Pace et al., 2018).

Sensitization of TRPV1 can be mimicked *in vitro* by incubating cultured sensory neurons with any of the aforementioned inflammatory mediators. In this experiment, the sensitizing agent is applied in between puffs of low doses of capsaicin (Bonnington and McNaughton, 2003). An increase in the magnitude of the calcium influx to the subsequent capsaicin stimulus –compared to the capsaicin response before incubation with the inflammatory mediator– is indicative of TRPV1 sensitization. We tested the ability of the TRPV1- expressing NOCL3 cells to become sensitized by the inflammatory mediators NGF and Bradykinin as well as the phorbol ester PMA. We found that hESC-derived nociceptors can be sensitized by the PKC activator PMA and the cognate TRKA ligand NGF, while Bradykinin was not able to sensitize the cells (Fig 5B–D). This sensitization profile was in agreement with the expression profile of the respective inflammatory receptors determined by transcriptional profiling: while TRKA was present in NOCL3 neurons, the Bradykinin receptor was not detectable (Fig 3C).

Collectively, we conclude that transient expression of *NGN1* in hESC-derived NCLCs, cultured in a defined serum-free medium, results in the generation of a molecularly homogenous population of *TRKA*- and TRPV1-positive population that displays typical features of inflammation-responsive nociceptors.

### Electrophysiological properties of human nociceptor-like cells

We confirmed that all derived nociceptor types, NOCL1 to NOCL4, are excitable and exhibit action potentials (APs) upon current injection. Overall, AP half widths of NOCL1-3 preparations were similar and well in the range of native intermediate-size murine nociceptors that typically give rise to Aδ fibers (Fang et al., 2005; Lechner et al., 2009; Rose et al., 1986) (Fig 6A). AP waveforms of cells derived according to the NOCL4 protocol, showed a broader AP width, including a ‘shoulder’ in the AP decay phase (Fig 6A, Fig EV7A) –a typical ion current feature found in some human and mouse nociceptors (Davidson et al., 2014). Additionally, we recorded resting membrane potentials in the range characteristic for small-diameter (C-fiber-type) human and mouse DRG neurons (Fang et al., 2005) (Fig EV7B).

**Figure 6.**
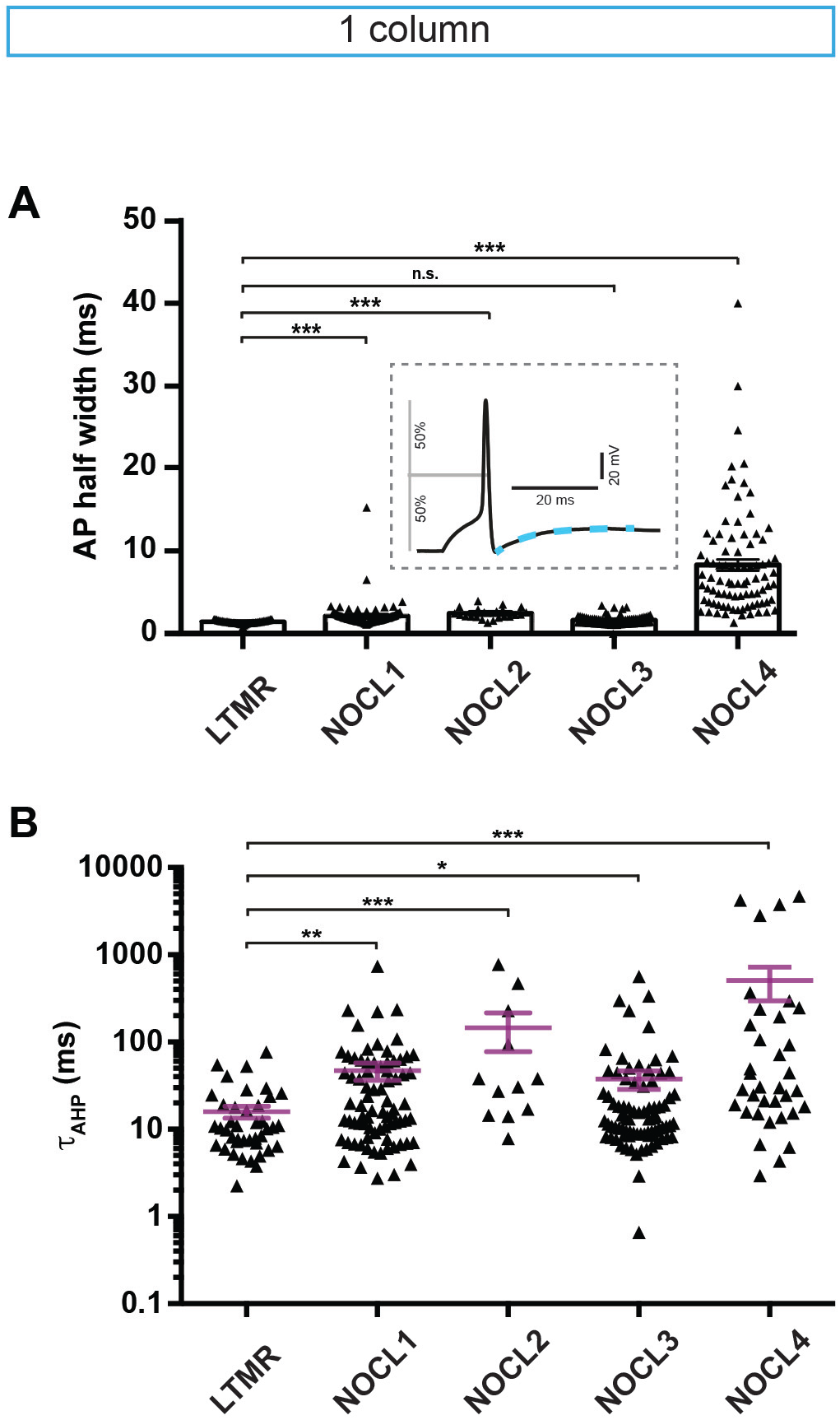
Electrophysiological characterization. A. Somatic action potential (AP) half width determined as indicated in the example shown in the inset; LTMR (n= 43), NOCL1 (n=109), NOCL2 (n=70), NOCL3 (n=90) and NOCL4 (n=87) were compared by Mann-Whitney test, n.s. (p=0.1243), ** (p=0,0023), *** (p<0.0001). B. Recovery from after-hyperpolarization was fitted with a single exponential function (blue dashed line in (A). n.s. (p=0.1522), * (p=0.0337), ** (p=0.0020), *** (p<=0.0003), Mann Whitney test. In (A) and (B) each triangle in the scatter plots corresponds to one cell, mean ± SEM is depicted.

These electrical attributes correlated well with estimated cell sizes assessed by capacitance measurements: The size range of NOCL1-3 neurons were comparable to small/intermediate-sized native human nociceptive neurons (Davidson et al., 2014; Rostock et al., 2018). On average, NOCL4 neurons turned out to be slightly smaller (Fig EV7C). The average size of hES-derived mechanoreceptors (Schrenk-Siemens et al., 2015), was larger than that determined for NOCL1-4 neurons, in agreement with smaller soma sizes found in native human nociceptors compared to LTMRs (Rostock et al. 2018).

Of note, continuous expression of *NGN1* (NOCL2 and NOCL4) led to smaller cell sizes compared to transient *NGN1* expression (NOCL1 and NOCL3) (Fig EV7C).

Extent and duration of after-hyperpolarizing currents are a defining feature of nociceptors and underlying ion channels of the HCN family. These channels have been implicated in pain-signal transmission (Emery et al., 2012; Tsantoulas et al., 2017). All four differentiation procedures (NOCL1-4) showed a prolonged rise time constant following the after-hyperpolarization potential (Fig 6B), similar to those reported for human and mouse nociceptors (Davidson et al., 2014; Momin et al., 2008; Schrenk-Siemens et al., 2015). Interestingly, the average decay time of hES-derived LTMRs was shorter than those recorded in any of the four NOCL subpopulations. This correlates with *HCN1* expression in the LTMRS (Fig 3C), a HCN variant largely absent from derived NOCL neurons and native mouse nociceptors (Viana and Belmonte, 2008). Conversely, NOCL neurons, in particularly NOCL3s, expressed *HCN2* (Fig 3C), an ion channel associated with pathological forms of pain (Momin et al., 2008; Tsantoulas et al., 2017).

Human and mouse nociceptors express both TTX-(Tetrodotoxin-) sensitive and TTX-insensitive voltage-gated sodium (Na_V_-) channels. Both channel types have been implicated in pathological forms of pain in human patients and mouse models (Habib et al., 2015). Accordingly, we found both types of conductances present in NOCL neurons. Irrespective of the differentiation protocol utilized, a small fraction (around 5-10 %) of sodium currents was TTX resistant, similar to currents recorded in a subset of native murine small diameter DRG neurons that we sampled randomly (Fig EV7D). Current-voltage relationships for NOCL3 in the presence of 300-500 nM TTX are indicative of a substantial contribution of Nav 1.8 currents, a marker of nociceptors (Fig EV7E) (Inserra et al., 2017). Nav 1.8 contribution is further enhanced over time (≥ 24 days vs. ≥ 48 days in culture; Fig EV7E inset).

In summary, characteristic ionic current properties of native nociceptors are recapitulated in derived NOCL neurons, with subtle variations observed across the 4 different differentiation procedures. Together, differences between NOCL1-4 neurons therefore resemble heterogeneity found in native DRG nociceptors.

### Investigating the role of PIEZO2 in hESC-derived and native mouse nociceptors

Many nociceptors are polymodal neurons that are able to respond to different stimuli such as temperature, acid, pungent chemicals and mechanical force. Other nociceptors are more specifically tuned to respond to discrete stimuli. We found NOCL3 neurons to respond to noxious temperature as well as pungent substances such as capsaicin, while only a small fraction of NOCL1- and NOCL2 neurons responded to capsaicin (Fig 2B–C). This was also reflected in the higher *TRPV1*-exoression in NOCL3 cells (Fig 3C).

Next, we investigated whether derived nociceptors could respond to mechanical stimulation. Differentiated neurons were mechanically stimulated by indenting the somatic cell membrane using a nanomotor driven probe, while simultaneously recording somatic currents –a typical stimulation paradigm that recapitulates mechanosensitivity of native DRG sensory neurons (Coste et al., 2010; Schrenk-Siemens et al., 2015). Indentation of the cell membrane in steps of 1 µm resulted in increasing inward currents in many NOCL neurons, showing their ability to respond to mechanical stimulation (Fig 7A and Fig EV7E).

**Figure 7.**
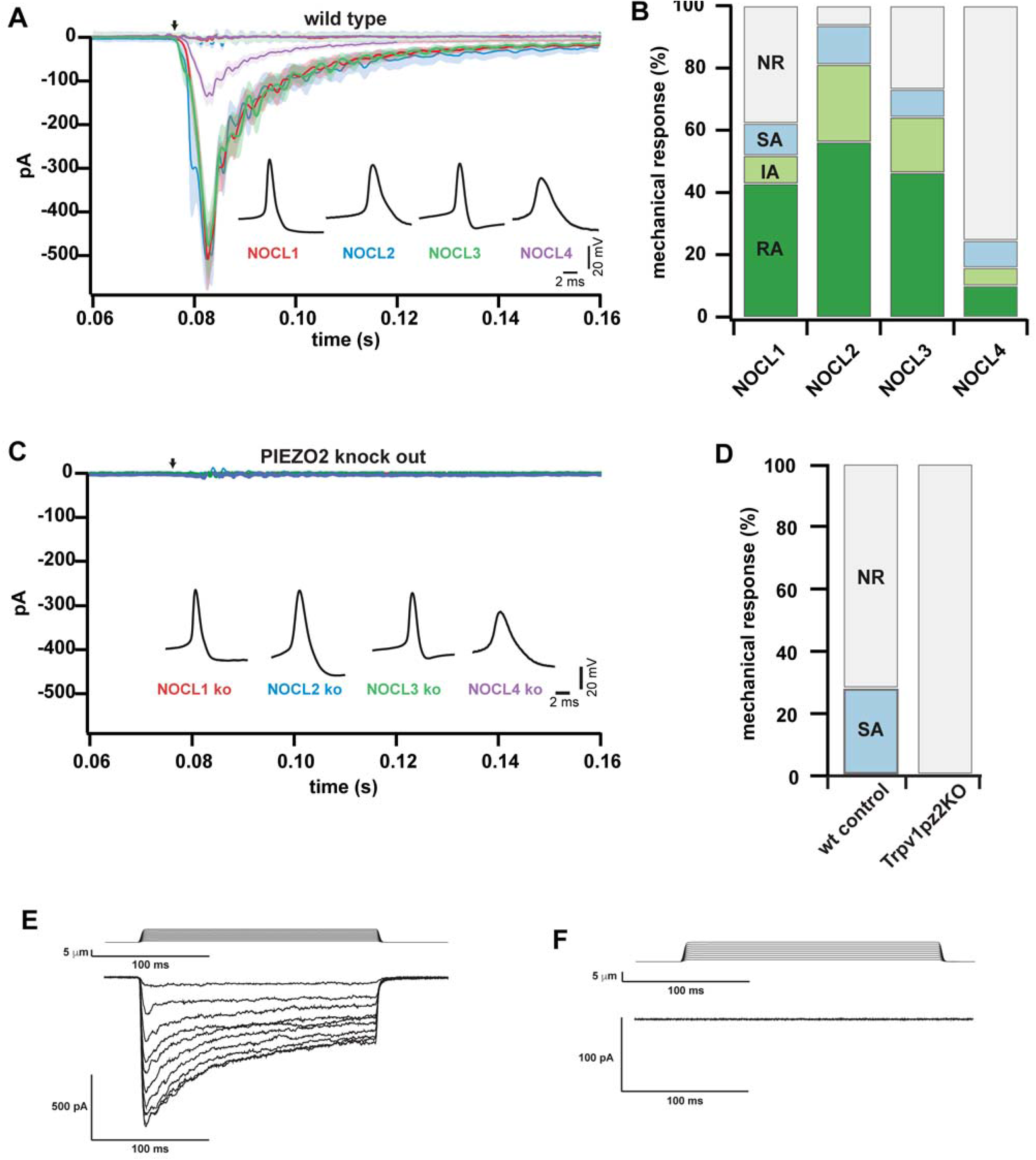
Mechanosensitivity of NOCLs and mouse DRG neurons. A. Average currents (± SEM) of wild type cells, evoked by a mechanical somatic displacement of 10 µm (arrow) are shown. Insets show example APs recorded from respective NOCLs. NOCL1 (n=48 responders, n=29 non-responders), NOCL2 (n=38 responders, n=11 non-responder), NOCL3 (n=41 responders, n=15 non-responders), NOCL4 (n=17 responders, n=52 non-responders). B. Stacked bar charts show the percentage of neurons with rapidly-adapting (RA), intermediate-adapting (IA) or slowly-adapting (SA) mechanical current as well as non-responders (NR). C. Average currents (± SEM) of PIEZO2 KO neurons, evoked by a mechanical somatic displacement of 10 µm (arrow) are shown. Insets show example APs recorded from respective NOCLs. NOCL1 (n=21), NOCL2 (n=48), NOCL3 (n=29), NOCL4 (n=32). D. Stacked bar charts show the percentage of mouse TRPV1/PIEZO2 wild type or TRPV1 wild type PIEZO2 knockout DRG neurons responding to mechanical stimulation with a slowly-adapting (SA) current or no current at all (NR). E and F. Currents of TRPV1/PIEZO2 wild type (E) and TRPV1 wild type PIEZO2 knockout DRG neurons evoked by increasing mechanical somatic displacement.

Classically, native murine mechanosensitive currents are classified according to their inactivation kinetics into rapidly, intermediately and slowly inactivating currents (Ranade et al., 2014). While LTMRs typically inactivate rapidly, nociceptors are markedly heterogeneous and comprise cells displaying all three types of inactivation categories (Viatchenko-Karpinski and Gu, 2016). Indeed, we were able to record all three types of mechanical current categories in the three NOCL preparations (Fig 7B).

NOCL1, NOCL2 and NOCL3 neurons had comparable fractions of mechanosensitive neurons, while the NOCL4 procedure resulted in less neurons responding to membrane indentation (Fig EV7E).

PIEZO2 has been identified as a primary mechanical transducer in non-nociceptive LTMRs (Ranade et al., 2014; Schrenk-Siemens et al., 2015). However, both in murine and human nociceptors its contribution to mechano-nociception remains controversial (Murthy et al., 2018; Szczot et al., 2018). We recently generated a hES cell line devoid of PIEZO2 (Schrenk-Siemens et al., 2015). Using this cell line, we first tested whether knockout hES cells were able to generate NOCL neurons. We found that their potential to differentiate was indistinguishable compared to that of wild type control cells (Fig EV8A–B).

We then tested whether PIEZO2 would contribute to mechanosensation. When comparing PIEZO2 knockout NOCL neurons with wild type controls we found that no mechanically-inducible current remained, irrespective of the differentiation protocol used, while general electric excitability remained unaltered (Fig 7C and Fig EV8C–D).

In agreement with the expression of PIEZO2 in native murine and human nociceptors (Murthy et al., 2018; Prato et al., 2017; Rostock et al., 2018; Szczot et al., 2018; Szczot et al., 2017) and with the expression of robust RNA of the mechanotransducer *PIEZO2*, all 4 different NOCL neuron types (Fig 3C and Fig 4C), resemble different subtypes of human nociceptors. These data suggest that PIEZO2 is a mechanotransducer in human-derived nociceptors. This highly penetrant phenotype was surprising, given that PIEZO2 appears to play little —if any— role in transducing painful mechanical stimuli in mice and man (Murthy et al., 2018; Szczot et al., 2018). We therefore wondered whether PIEZO2 is also required in native mouse nociceptors to mediate membrane indentation-induced mechanical responses. Given that TRPV1 is a prototypical marker for nociceptive neurons that we found to be co-expressed with PIEZO2 in NOCL3 cells, we focused on TRPV1-expressing mouse nociceptors. To identify TRPV1 expressing sensory neurons, a Cre-dependent tdtomato expressing reporter virus was introduced into the dissociated sensory neuron cultures of Trpv1cre and Trpv1cre::Piezo2fl/fl animals. 36 hours after viral transduction, the tdtomato expressing neurons were analyzed electrophysiologically for the presence of mechanically-induced currents. We found 27,8% of the TRPV1-expressing cells (5 out of 18) to respond to mechanical stimulation with a slow adapting current (Fig 7D–E). This is in agreement with previous data, reporting *Piezo2* being expressed in 24% of TRPV1 positive neurons (Coste et al., 2010). In the absence of PIEZO2 in the TRPV1 positive neurons, none of the stimulated cells (n=21) showed responses to mechanical stimuli (Fig 7D, F), supporting our finding, that PIEZO2 is indispensable for mechanical transmission in a specific subset of mechano-nociceptive sensory neurons in culture.

## Discussion

Using embryonic stem cell-derived neural crest precursors in combination with viral gene expression, we here describe the generation of human nociceptor-like cells. The molecular make-up and receptor armament of the generated sensory neurons varied as a function of (i) the duration of *NEUROGENIN1* expression and (ii) the availability of serum and/or neurotrophic factors in the culture medium.

One striking feature of the system is the induction of a differentiation program that –when combined with a positive selection procedure– allowed the production of a homogenous population of sensory neurons that display functional hallmarks of prototypical nociceptors. One protocol in particular, resulted in the generation of a homogenous population of TRPV1-expressing nociceptor-like cells as was assessed by functional Ca-imaging as well as *in situ* hybridization experiments. Additionally, the cells showed the ability to become sensitized by inflammatory mediators such as NGF. Furthermore, the derived nociceptor-like cells responded to temperature stimulation, another characteristic feature of TRPV1-expressing nociceptors (Caterina et al., 2000). In comparison to cultured mouse DRG neurons, the homogenous response of the human ESC-derived neurons is the most striking asset, showing the potential of the cellular system as a model for studying specific sensory-neuron subpopulations. This has, to the best of our knowledge, never been achieved by previous protocols that produced rather low yields of heterogeneous nociceptor-like cells or a different subpopulation (Blanchard et al., 2015; Chambers et al., 2012; Wainger et al., 2015). The added benefits described here are the result of a two-step differentiation protocol that first generates neural crest-like cells, the natural precursors of sensory neurons, and secondly differentiates them further into sensory neurons. Recapitulating this developmental intermediate (neural crest cells), rather than attempting to trans-differentiate fibroblasts or hES/iPS cells into nociceptors directly (Blanchard et al., 2015; Wainger et al., 2015), promotes more efficient sensory neuron generation.

Previous cellular classifications, largely based on work in rodents, have used marker gene expression profiles to categorize subgroups of nociceptors (Basbaum et al., 2009). The advent of single cell RNAseq methods has broadened this type of molecular classification and has highlighted ever smaller and more refined subgroups of sensory neurons (Li et al., 2016; Usoskin et al., 2015). Unfortunately, such molecular information is much more scarce for human nociceptors, largely due to DRG inaccessibility and the difficulty to obtain native human tissue samples. Nonetheless, molecular differences between human and rodent sensory neurons have been observed such as the more widespread overlap of the two neurotrophic receptors *TRKA* and *RET* in mature human nociceptors (Rostock et al., 2018), a feature that we also find in the derived NOCL neurons. Notwithstanding, one limitation of our approach is that it is difficult to unambiguously determine what precise *native* nociceptor subtype the respective NOCL neurons correspond to. Two factors contribute to this: first, the limited available molecular information on native human nociceptors precludes comparison at a similar detail and depth possible for rodent cellular systems. And second, our *in vitro* differentiation procedure does not fully recapitulate *in vivo* sensory neuron differentiation, which involves temporally- and spatially refined cues as well as target organ innervation, which shape sensory subtype specification.

Nevertheless, we compared derived NOCL neurons with previously generated rodent molecular nociceptor profiles and found striking similarities with afore defined subgroups: based on expression profiles deduced from different sensory neuron subtypes (Li et al., 2016; Usoskin et al., 2015) NOCL1 neurons rather resemble peptidergic neurons, while the marker expression pattern of NOCL3 neurons showed more overlap with those of murine non-peptidergic neurons.

The heterogeneity of endogenous nociceptors is also reflected in their functional properties, such that certain subgroups of sensory neurons are tuned to detect heat-, cold-, chemical- and/or mechanical stimuli (Le Pichon and Chesler, 2014). Particularly important pathologically (and difficult to treat therapeutically) are mechano-nociceptive characteristics: the ability to respond to noxious (and thus painful) mechanical forces. We investigated the mechanotransduction properties of our derived nociceptor populations and found most of them to be responsive to mechanical stimulation.

A major unresolved question pertains to the molecular mechanism(s) that enable nociceptive neurons to detect and respond to painful mechanical stimuli. One protein, PIEZO2, has recently gained prominent attention because it was found to be the main transducer of innocuous mechanical stimuli (such as touch) in non-nociceptive LTMRs, both in mice and men (Chesler et al., 2016; Ranade et al., 2014; Schrenk-Siemens et al., 2015). Concurrently, the gene was also found to be expressed in a wide range of human and mouse nociceptors (of both, peptidergic and non-peptidergic origin), raising the eminent possibility that the receptor is also required for mechano-nociception. *In vitro* experiments by Dubin et al. (Dubin et al., 2012) using cultured mouse DRG neurons showed an increase in PIEZO2-dependent current amplitude and a slowing of its inactivation upon stimulating inflammatory pathways, suggestive of a role for PIEZO2 in inflammatory pain. Another study demonstrated a connection between capsaicin induced TRPV1 activation and subsequent PIEZO2 inhibition in nociceptors based on the depletion of membrane phosphoinositides (Borbiro et al., 2015). Additionally, in *d. melanogaster*, the PIEZO2 analog *dpiezo* is required for mechano-nociception but not for innocuous gentle touch (Kim et al., 2012). While these previous studies argue in favor of a role for PIEZO2 in mechano-nociception, more recent *in vivo* studies suggest that PIEZO may only play a minor –if any– role for this highly medically relevant property. One study, genetically deleting PIEZO2 in selective neurons in mice, suggested that the mechanotransducer plays a minor role in detecting painful stimuli such as pinch or pinprick (Murthy et al., 2018). In another study, focusing on 4 human patients with point mutations in the *PIEZO2* gene, a contribution for a function of the protein in acute mechano-nociception could not be found (Szczot et al., 2018).

To our surprise and in contrast to what has been reported from the aforementioned patient studies, absence of PIEZO2 in the hESC-derived nociceptor-like cells, using a previously generated PIEZO2 knockout hESC line (Schrenk-Siemens et al., 2015), showed the loss of mechanotransduction in all generated subpopulations. We found a similar result in cultured mouse nociceptors, where about 27% of TRPV1 expressing mouse neurons showed mechanotransduction properties, while none could be detected in the absence of PIEZO2.

The discrepancy between the patient studies and our results could be partly due to the mechanical stimulation paradigm used in cell culture to mimic physiological mechanical pain stimuli: high frequency indentation of the plasma membrane using a nanomotor-driven probe. While our results suggests that there is a requirement for PIEZO2 in indentation-mediated mechanotransduction at the cell soma in both, human ESC-derived nociceptor-like cells as well as in a subset of primary mouse TRPV1-expressing DRG neurons, additional mechanisms may contribute to mechanosensitivity under *in vivo* conditions for example at (skin-) innervating fiber terminals.

Another possible explanation could be a potential contribution of other mechanosensitive ion channels in nociceptors of patients devoid of functional PIEZO2. Several other ion channels with mechanosensitive properties have been described in DRG neurons (Gu and Gu, 2014) and their mechanosensitive properties may become unmasked to play more prominent roles in the absence of PIEZO2.

In summary, our differentiation protocols allow for the generation of several different nociceptor subtypes that bear prominent molecular and functional similarity to rodent nociceptor subgroups. Furthermore, our cellular system offers the possibility of overcoming the difficulties associated with lethality and/or developmental compensation of loss-of-function mutations that are often observed in mice or other model organisms. By recapitulating nociceptor development *in vitro*, this cellular system can help to unravel unrecognized functions of sensory signaling pathways and molecules (such as PIEZO2), thereby complementing work in more complex organisms.

## Material and Methods

### Generation of hESC-derived sensory neurons

#### 1) Generation of hESC-derived neural crest-like cells (NCLCs; protocol adapted from Bajpai et al., 2010)

Use of Human embryonic stem cells (HUES-7 line; Harvard University, Cambridge, MA, USA) was approved by the German Central Ethics Committee for Stem Cell Research (project codes: #1710-79-1-4-51 and 1710-79-1-4-51- E01). hESCs were cultured on Matrigel (Corning) coated dishes in E8-medium (self-made according to Chen et al., 2011) at 37°C and 5% CO_2_ until 80% confluency. Cells were detached using EDTA (0.5 mM) and transferred on uncoated dishes in sphere medium (DMEM:F12, Neurobasal, N2-supplement, B27-supplement, Glutamine (all ThermoFisher Scientific), Insulin (CSBio) with freshly added EGF and FGF (20 ng/μl; Peprotech). Rock inhibitor (10 μM; Stem Cell Technologies) was added for up to 48 h, depending on size of spheres. One day after plating, small spheres become visible. Medium was changed every other day. After 7-9 days, spheres spontaneously attach and NCLCs start to migrate out. NCLCs were collected using Accutase (Sigma) after remnants of spheres were manually removed from the plate. NCLCs can be frozen in liquid nitrogen or immediately plated on Poly-D-ornithine (15 μg/μl, Sigma), laminin (10 μg/ml, Sigma) and fibronectin (10 μg/ml, ThermoFisher Scientific) coated 3.5 cm dishes in a density of 60000 cells/cm^2^ in sphere medium without FGF and EGF.

#### 2) Generation of low threshold mechanoreceptors (LTMRs)

LTMRs were generated as described in Schrenk-Siemens et al., 2015. In brief: NCLCs were cultured in differentiation medium (sphere medium supplemented with BDNF, GDNF, NGF and NT3 (all 10 ng/μl, Peprotech) and retinoic acid (0.1 μM, Sigma). Medium was changed daily and every second day starting after two weeks. LTMRs were used for experiments after at least 21 days in culture.

#### 3) Generation of nociceptor-like cells (NOCL1-3) using induced *NGN1* expression

To generated NOCL1-3, NCLCs were infected with two lentiviruses: one expressing rtTA under the control of an ubiquitin promoter for constitutive expression of rtTA. The other expressing *NGN1*, *EGFP*, and a puromycin resistance cassette as a fusion protein, linked via P2A and T2A sequences, under the control of TeTO promoter. Cells were infected in sphere medium, containing HEPES (10 mM, ThermoFisher Scientific) and protamine sulphate (8 μg/ml). After 6 h, cells were washed three times with PBS and kept in sphere medium until the next day.

The next day, medium was changed to differentiation medium (sphere medium containing BDNF, NGF and GDNF (10ng/ml; Peprotech), Antibiotic-Antimycotic (1x; ThermoFisher Scientific) and in the case of NOCL1 and NOCL2 also 5% fetal calf serum (ThermoFisher Scientific). *NGN1* expression was induced by addition of doxycycline (2 μg/ml, Sigma) for 10 days (NOCL1 and 3) or for the whole culture period (NOCL2). On the second day of induction, puromycin (10 μg/ml, Invivogen) was added (NOCL3) for 24-48h to eliminate those cells that were not infected by the virus. After 4 to 5 days of induction, cells were detached using Accutase (Sigma) and plated on freshly laminin/fibronectin coated dishes or glass coverslips in a density of 31000 cells/cm^2^. After 13 days in culture, cells were treated with Mitomycin C (10 μg/ml, Sigma) for 45 min, to prevent proliferation of any non-neuronal cells still in the culture, then washed carefully twice with PBS and kept in differentiation medium for at least 21 days in total. Medium was changed every day for the first 10 days and then only every other day.

#### 4) Generation of nociceptor-like cells using *NGN1*, *ISL2* and *KLF7* (NOCL4)

Modified from the protocol published by Wainger et al., 2015, NOCL4 cells were generated by infecting NCLCs with three lentiviruses expressing constitutively *NGN1*, *ISL2* and *KLF7*. Cells were infected for 6 h, then washed with PBS and kept in sphere medium until the next day. Medium was then changed to differentiation medium (as above) including 5% fetal calf serum (ThermoFisher Scientific). Neurons were kept in culture for at least 21 days before experiments were done.

### Virus Generation

Lentiviruses were produced in HEK293TN cells (Biocat) by cotransfection with three helper plasmids (pRSV-REV, pMDLg/pRRE and pMD2.G, obtained from Dr. Alexander Löwer, Berlin, Germany) plus the lentiviral vector DNA: FUW-TetO-Ngn1-P2A-EGFP-T2A-Puro, FUW-rtTA (obtained and modified from Dr. Thomas Südhof, Stanford, USA); pMXs-dest-WRE-Ngn1, pMXs-dest-WRE-Klf7, pMXs-dest-WRE-Isl2 (obtained from Dr. Clifford Woolf, Harvard Medical School, MA, USA); ELFa-Runx1-IRES-GFP, ELFa-TRKA-IRES-GFP using calcium phosphate transfection. Lentivirus containing medium was harvested 48 and 72h after transfection, centrifuged (800 x g for 20 min) and filtered (0.45 µm pore size). The virus was subsequently concentrated by mixing the filtered supernatant with 50% PEG-8000 (Sigma) in a ratio of 4:1 and placed at 4°C over night. The concentrated viral particles were pelleted by two centrifugation steps (2,500 rpm for 20 min, followed by 1,200 rpm for 5 min), resuspended in 200 µl of PBS and stored at −80°C.

### Generation of *TRKA*-tomato reporter hESC-line

A *TRKA*-tomato reporter hESC line was generated using homologue recombination with the help of CRISPR/Cas9 technology. To this end, a targeting construct was cloned containing a tandem tomato sequence linked via a T2A sequence to the 5’ homology arm containing parts of *TRKA*-Exon 17, as well as to a 3’ UTR site. The vector also included a blasticidin selection cassette. *TRKA* specific target sites for CRISPR-Cas9 were designed using the CRISPR design online tool from either DNA2.0 (now ATUM) or from the Jack Lin CRISPR/Cas9 gRNA finder. Two oligos were used: oligonucleotides score 1: CACCGCTGGGCTAGGGGGCCGGCCC-forw; AAACGGGCCGGCCCCCTAGCCCAGC-rev and oligonucleotides score 2: CACCGCCTGGATGTCCTGGGCTA-for; AAACTAGCCCAGGACATCCAGGC-rev, were cloned into CRISPR:hSpCas9 (px330, Adgene) and used for nucleofection. To this end, a single cell suspension of hESCs was prepared using EDTA and thorough pipetting. Cells were spun at 1000 rpm and counted. 8 x 10^5^ cells were resuspended in 100 μl Ingenio-solution (Mirrus) supplemented with Rock inhibitor (10 μM, Y-27632, Stem Cell Technologies) and 2.5 μg of CRISPR:hSpCas9 plasmid and 2.5 μg linearized DNA targeting plasmid and immediately pulse with program DC100 (using a 4D-Nucleofector, Lonza). Cells were resuspended in warm mTeSR1 (Stem cell technologies) supplemented with Rock-inhibitor (10 μM) and plated on one Matrigel coated 10 cm dish. Blasticidin selection (2 μg/ml, InvivoGen) was started 2–4 days after plating of cells and picking of clones was started 10–14 days after nucleofection. Picked clones were plated on Matrigel coated 96-well plates in the presence of Rock-inhibitor. Rock inhibitor was left out 1–2 days after picking. For Southern blot analysis, DNA was isolated from 96 well plates. Several positive clones for *TRKA*-tomato hESCs were derived and tested for their differentiation potential. The same strategy was used to generate a *TRKA*-tomato *PIEZO2^−/−^* hESC line, using the previously generated *PIEZO2^−/−^* hESC line^2^ as starting material.

### Primary cultures of DRG neurons

All animal work was approved by the Ethics and Animal Welfare Committee of the University of Heidelberg and the State of Baden-Wuerttemberg. C57BL/6 female mice at 6 weeks of age were sacrificed using isoflurane and subsequent decapitation. DRGs from all spinal segments were collected in Ringer solution. DRGs were subsequently treated with collagenase IV (1 mg/ml; Sigma-Aldrich) and trypsin (0.05%, Invitrogen) at 37°C for 60 min and 15 min, respectively, at 37°C. After enzymatic treatment, DRG’s were washed twice with growth medium [DMEM-F12 (ThermoFisher Scientific) supplemented with L-glutamine (2 µM; Sigma-Aldrich), Antibiotic-Antimycotic (1x; ThermoFisher Scientific) and 5% fetal calf serum (Invitrogen)], triturated using a 1 ml pipette tip and plated in a droplet of growth medium on a glass coverslip precoated with poly-L-lysin (20 µg/cm²; Sigma-Aldrich) and laminin (4 µg/cm²; Invitrogen). To allow neurons to adhere, coverslips were kept for 1 h at 37°C in a humidified incubator before being flooded with fresh growth medium. Cultures were used for Ca-imaging experiments on the 3rd day after preparation.

### Primary cultures of DRG neurons from wild type and Piezo2 knockout mice

Trpv1-Cre::piezo2fl/fl were bred by crossing the Piezo2^fl/fl^ (Szczot et al., 2018) with Trpv1-IRES-Cre animals (Cavanaugh et al., 2011). Dorsal root ganglia (DRGs) cultures were made from adult Piezo2^fl/fl^:: Trpv1-IRES-Cre or Trpv1-IRES-Cre mice. Animals were euthanized and DRGs harvested in accordance to the National Institutes of Health (NIH) guidelines approved by the National Institute of Neurological Disorders and Stroke (NINDS) Animal Care and Use Committee. For enzymatic dissociation the ganglia were incubated and gently agitated at 37°C in collagenase (1mg/ml in PBS, Sigma-Aldrich St. Louis, USA) for 5-20 minutes. The collagenase solution was then removed and replaced with Trypsin (0.25%, ThermoFisher, Waltham, USA) for 2-10 minutes. Following the enzymatic treatment, the Ganglia were triturated with a fire polished Pasteur pipette in culture media (88% MEM (Gibco, Gaithesburg, USA), 10% Fetal Horse Serum (Gibco), 1% Pen/Strep (Lonza, Basel, Switzerland), 1% Vitamin mix (Gibco)). The supernatant was then transferred to a 15 ml tube (Corning, Tewksbury, USA) containing 6ml of culture media and centrifuged for 10 minutes at 150g to concentrate the dissociated sensory neurons into a pellet. The supernatant was removed and the pellet was resuspended in a mixture of 100ul of culture media and 5ul of a reporter virus (pAAV-CAG-LSL-tdtomato (cat no. 100048, University of Pennsylvania viral core, USA)). A 10 ul drop of the cell suspension was then plated onto Poly L-Lysine/Laminin coated coverslips (NeuVitro, Vancouver, USA) in 24 well-plates (Corning). The plates were placed in an incubator (37°C/5% CO_2_) and, after an hour of incubation, 500ul of the culture media was added into each well. Coverslips of sensory neurons were used for whole cell recording experiments 36-72 hours after plating.

### Immunocytochemistry

Cells were fixed in 4% PFA for 10 min at room temperature and washed three times with PBS, before blocking solution (10% goat serum (PAN Biotech) in PBS) was added for 1 h at room temperature. Cells were incubated in primary antibodies solution (antibody diluted in PBS containing 3% goat serum, 0.01% Triton x-100 (Merck)) at 4°C over night. Cells were washed three times with PBS and secondary antibodies and DAPI (1:12000) were added to the cells in PBS containing 3% goat serum and 0.01% Triton X-100 for 2 h at room temperature in the dark. Cells were washed three times with PBS and mounted on glass slides with self-made mowiol. Stainings were dried at room temperature over night in the dark and kept on 4°C further on.

Primary antibodies used:

Chicken anti Neurofilament 200 (1:25000; Abcam, #ab72996), mouse anti ISL1 (1:100; Developmental Studies Hybridoma Bank, #39.4D5), rabbit anti MAFA (1:20000; obtained from Dr. Carmen Birchmeier, Berlin, Germany). Alexa 488-, 555- and 647-conjugated secondary antibodies were obtained from Molecular probes and used 1:1000.

### *In situ* hybridization combined with immunocytochemistry

*In situ* hybridization was carried out as previously described (Wende et al., 2012) and optimized for human cells. HESC-derived neurons were fixed for 30 min in 4% PFA, acetylated and permeabilized in 0.3% TritonX–100 in PBS. After pre-hybridization, hydrolyzed DIG- and/or FITC-labelled RNA probes were added and hybridization was performed over night at 60°C. The next day cells were washed twice with 2 x SSC/50% Formamide/0.1% N-Lauroylsarcosine at 60°C and then treated with 20 µg/ml RNase A for 15 min at 37°C. After washing twice in 2 x SSC/0.1% N-Lauroylsarcosine and twice in 0.2 x SSC/0.1% N-Lauroylsarcosine at 37°C for 20 min each, cells were blocked in MABT/10% goat serum/1% blocking reagent.

For a double fluorescent *in situ* hybridization, sections were stained by two sequentially rounds of Thyramide signal amplification (TSA) steps with an intermediate peroxidase inactivation with 3% H_2_O_2_ for 2 h and 4% PFA for 30 min. Tissue was incubated with anti-FITC-POD (1:2000; Roche) or anti-DIG-POD (1:1000; Roche) over night at 4°C. After washing in MABT, the TSA reaction was performed by applying either Thyramide-Biotin on the third or Thyramide-Rhodamine on the fourth day for 30 min at room temperature. For the detection of Thyramide-Biotin a streptavidin-cy2 antibody (1:1000; dianova) was applied, whereas nuclei were stained with DAPI. Subsequently cells were washed and blocked in 10% NGS in PBS for 1 h, before an immunostaining was started as decribed above.

*In situ* probes for *PIEZO2* and *RET* were obtained from Dr. Hagen Wende (Institute of Pharmacology, University of Heidelberg). Full length clones for human *TRPV1* and *TRPA1* were provided by Dr. David Julius (University of California, San Francisco), and the generated *in situ* probes covered the complete open reading frame. The remaining *in situ* probes were amplified using the following primers and cloned into pBluescript SK(+).

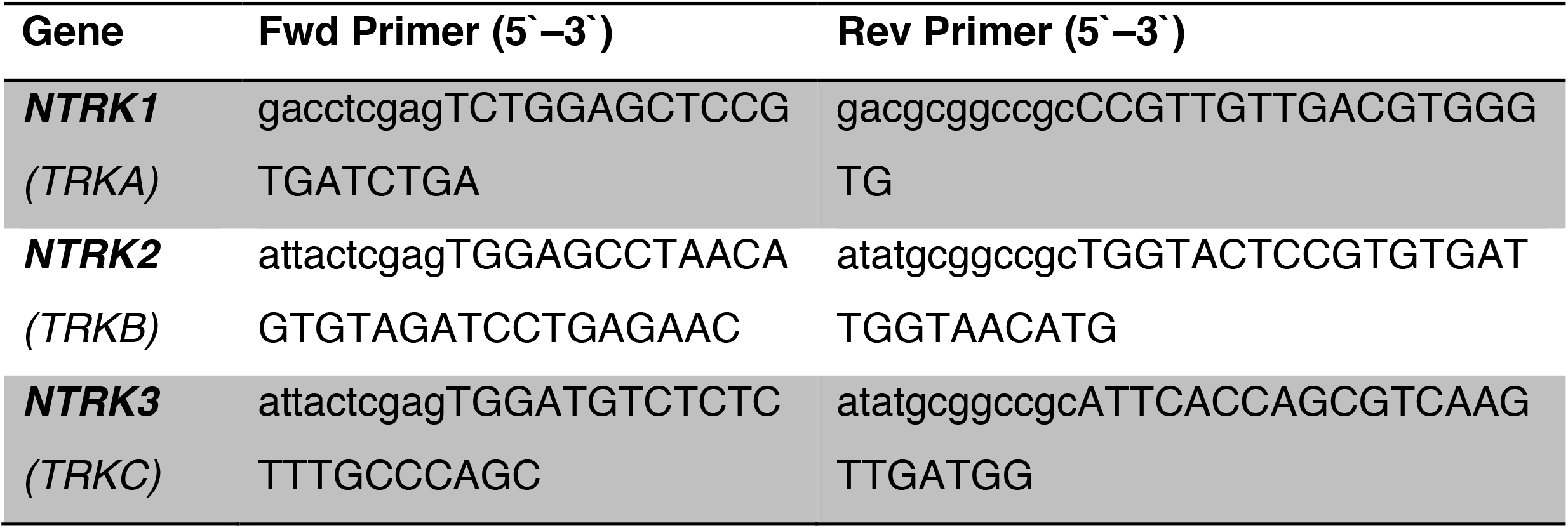

### Imaging and quantification

Fluorescence images were taken on a Zeiss Axio Observer A.1 inverted microscope using a Cool Snap camera from Visitron Systems or a Nikon Eclipse Ts2 microscope.

In situs and immunocytochemistry for the indicated markers were performed on at least three independent differentiations. For quantification, cells were stained after indicated times in culture and random pictures were taken from at least three different differentiations. Positive cells were counted using image J software.

### RNA-Seq Sample generation and Anaylsis

For LTMRs, RNA-sequencing data were taken from Schrenk-Siemens et al., 2015. For NOCL1 and NOCL2, clumps of 10–15 cells were picked individually and amplified using the Smart-Seq2 protocol (Picelli et al., 2014).

For NOCL3 RNA was isolated at different time points (0, 12, 24 and 31 days after induction of *NGN1*) from whole 3,5 cm dishes using Trizol reagent (Thermo Fisher scientific), based on Chomczynski and Sacchi, 1987. Integrity of the extracted RNA was investigated, using the Agilent RNA 6000 Bioanalyzer Nano Kit (nCounter Core Facility, Institute of human genetics, University Heidelberg). Libraries were generated using the Illumina TruSeq mRNA library prep kit. Samples were run on an Ilumina NextSeq 500 sequencer on paired-end mode with 75 bp (Genomics Core Facility, EMBL, Heidelberg). Fastq files were aligned using the GRCh38.p12 assembly transcriptome acquired from GENCODE and the Salmon pseudoalignment package (Patro et al., 2017). Quality control was performed using FastQC and MultiQC (Andrews 2010; Ewels et al., 2016). Alignemnts were imported into R using the tximport package and normalized counts calculated using DESeq2 (Soneson et al, 2015; Love et al., 2014).

### Ca^2+^-imaging

HESC-derived nociceptor-like cells, LTMRs or mouse DRG neurons were loaded with FURA-2 AM (10 μg/ml, HelloBio) or Cal520 AM (5 μM, AAT Bioquest) in Ringer [(mM): 140 NaCl, 5 KCl, 2 CaCl_2_, 2 MgCl_2_, 10 HEPES and 10 glucose, adjusted to pH 7.4] with pluronic acid, F-127 (0.5 μg/ml; ThermoFisher Scientific) for 1 h at room temperature or 37°C respectively, in the dark. Cells were washed twice with Ringer and kept at room temperature for at least 20 min before starting the experiments. Fluorescent images were acquired with Metaflour Software (Molecular Devices), traces extracted using Suite2p and analysed with image J and/or Graph Pad Prism Software. For standard experiments, cells were challenged with Capsaicin (1 μM, Tocris), Menthol (500 μM; Sigma) or Mustard Oil (Allyl-isothiocyanate, 200 μM; Sigma). For sensitisation experiments cells were challenged 5–times for 1 min with Capsaicin (10 or 100 nM), with 5min of washing with Ringer in between. After the 5^th^ stimulation, cells were incubated for 5 min with either NGF (nerve growth factor, 100 ng/μl; Peprotech), PMA (phorbol 12-myristate 13-acetate, 1 μM; Sigma), Bradykinin (10 nM; Tocris) or Serotonin (100 μM, Tocris), followed by another stimulus of Capsaicin (10 or 100 nM) for 1 min. To assess the amount of Capsaicin responders in general, another Capsaicin challenge (1 μM) was given after 5 min washing. For all Ca-imaging experiments, cells were challenged with high K^+^ Ringer solution [(mM): 45 NaCl, 100 KCl, 2 CaCl_2_, 2 MgCl_2_, 10 HEPES and 10 glucose, adjusted to pH 7.4] at the end to visualize all neurons in the culture. For analysis, the average fluorescence ratios (F_340_/F_380_) for capsaicin-only, mustard oil only, mustard oil and capsaicin or high potassium only were calculated and shown as traces or parts of whole tables. For the analysis of the sensitization experiments, the ratio of calcium responses between peaks 6 and 4 (second peak before and first peak after incubation with NGF, PMA, serotonin or bradykinin) was compared.

To test the temperature response of the NOCL3 in comparison to mouse DRGs, temperature challenges of 25 s duration were given to the cells, with 5 min intervals at room temperature in between. Different temperature stimuli were generated using a self-made glass coil system connected to a water bath, temperature at the cells was measured using a temperature probe (physitemp) and increased temperature steps (starting at 35°C, until 48°C) were used. Slight variations in the temperature at the side of the measurement could be observed due to thermal bridges in the system. To assess TRPV1 positive cells and neurons in general, a capsaicin stimulus (1 μM) and high potassium stimulation respectively, were given after the last temperature stimulus. An acquisition rate of 10Hz was used in combination with the calcium dye Cal520-AM. Cells were segmented and traces extracted using the Suite2p pipeline (Pachitariu). R was used to calculate the dF/F0 ratios that were then used to generate the heatmaps using the ComplexHeatmap package (Gu et al., 2016).

### Electrophysiological Recordings

#### Patch clamp recordings

Cell culture medium from cells cultured on 3,5 cm dishes was exchanged with ∼4 ml pre-warmed (37°C) external solution consisting of (in mM) 136 NaCl, 4 KCl, 2 CaCl_2_, 1 MgCl_2_, 10 D-glucose, 10 HEPES, adjusted with NaOH to pH 7.4, resulting in ∼295 mOsm. Using an inverted microscope (Zeiss, Axio Observer A1), differential interference contrast or phase contrast light images were acquired with a 20X objective. Red fluorescence was tested with a HXP 120 C light source and a mCherry filter set. Whole cell patch clamp recordings were performed with an Axopatch 200B amplifier, sampled at 20-100 kHz, low pass filtered at 10 kHz and digitized with a Digidata 1440A using Clampex 10.6. Recordings were performed at room temperature (22-24°C) up to 4-6 h with borosilicate glass (O.D. 1.5 mm, I.D. 0.86 mm, Sutter Instrument, BF150-86- 7.5), pulled on a micropipette puller (P-97, Sutter Instrument) with an open pipette resistance of ∼2-4 MΩ measured with a KCl- or CsCl-based patch pipette solution, subsequent gigaseal >1 GΩ. Patch pipettes were filled with (in mM) 1 EGTA, 10 HEPES, >115 KCl, 6 MgCl_2_, 6 Na_2_-phosphocreatine, 4 Na_2_- ATP, 0.3 Na-GTP, pH adjusted to 7.4 with KOH and osmolarity adjusted with further addition of KCl to 280-285 mOsm. Neurons were held at −65 mV in voltage clamp.

Cell capacitance was assessed in voltage clamp by hyperpolarizing the cell by 15 mV for 100 ms, averaging 25 response current traces and fitting a biexponential decay to baseline to the first capacitance decay. Cell capacitance was calculated by dividing the weighted biexponential decay by the cell’s input resistance. The cell’s input resistance was calculated by dividing the applied voltage step by the relative current of the first capacitive transient.

Action potentials were measured in current clamp with active bridge balance. Cells were held at −65 mV by injecting current. Increasing current steps of 100 ms or 1 s were applied to evoke action potentials.

The occurence of a ‚shoulder’ in the falling phase of the action potential was quantified by fitting a monoexponential function between the peak of the action potential and the minimum of its afterhypolarization (AHP), assessed as the voltage minimum within 20 ms after the peak of the action potential.

The rising of the after-hypolarization (τAHP) was assessed by a monoexponential fit of 32 ms length starting at the minimal voltage within 20 ms after the action potential peak, if the AHP amplitude was at least 0.1 mV and no second action potential was elicited.

Mechanically activated currents were recorded at −65 mV with a heat-polished glas pipette of the same properties as used for patch clamp, attached to a micromanipulator (Nanomotor, MM3A, Kleindieck Nanotechnik, Reutlingen, Germany) and the evoked whole cell currents were sampled at 20-100 kHz and hardware (Axopatch 200B) low-pass filtered at 10 kHz and subsequently software low-pass filtered with 1 kHz. The micromanipulator probe was positioned at an angle of 45° to the surface of the dish at opposing site to the patch pipette. The probe moved with a velocity of ∼1.3 µm/ms. Currents were fit by a bi-exponential function to the baseline before the stimulus and classified as rapidly-adapting (<10 ms), intermediate-adapting (10-30 ms) and slowly-adapting (>30 ms) mechanical currents according to their weighted biexponential inactivation time constant. Cells were classified as mechanically inactive when stimuli of <= 10 µm indentation did not elicit responses > −40 pA. Recordings in which leak currents significantly increased upon mechanical stimulation were not considered.

Amplitudes of mechanical evoked currents were either determined by the weighted amplitude of the biexponential fit or by measuring the relative amplitude.

For assessment of TTX-resistant sodium currents, a diamond-shaped recording chamber was constantly perfused (∼2 ml/min) with a gravity-driven ValveLink 8.2-system (AutoMate Scientific, USA) with (in mM) 128 NaCl, 20 TEA-Cl, 1 4-AP, 1 CaCl_2_, 1 MgSO_4_, 8 glucose, 10 HEPES, 0.1 CdCl_2_, adjusted to pH 7.4 with NaOH, resulting in ∼300 mOsm adding Tetrodotoxin citrate (HB1035, Hello Bio, Bristol, UK or cat. no. 1069, Tocris). Patch pipettes were filled with (in mM) 10 EGTA, 10 HEPES, >100 CsCl, 6 MgCl_2_, 4.8 CaCl_2_, 6 Na_2_- phosphocreatine, 4 Na_2_-ATP, 0.4 Na-GTP, pH was adjusted with CsOH to 7.4, osmolarity was adjusted by further addition of CsCl to ∼280-285 mOsm. In voltage clamp, with actively compensated pipette series resistance and whole cell capacitance (∼70%), cells were held at −65 mV, then hyperpolarized for 200 ms to −120 mV to recover TTX resistant currents, and then clamped for 40 ms to potentials from −50 mV to 50 mV in steps of 5 mV.

Analysis was carried out in Igor Pro 6.37 (WaveMetrics, USA), after importing Axon binary files with DataAccess (Bruxton Corporation, USA), or in Graphpad Prism 5 or 7 or 8. Statistical tests performed as indicated in the respective legends, were two-sided. Adjustment for multiple comparisons was not performed.

### Neuronal voltage clamp recording of mouse DRG cells

Whole-cell current responses of intersectionally targeted Trpv1 and tdtomato expressing neurons were recorded. Recordings were done using a Multiclamp 700B amplifier (Molecular Devices, Sunnyvale, USA) at a holding potential of −60mV. Signals were digitized with a Digidata 1550 (Molecular Devices) digitizer at 100 kHz, low pass filtered at 10 kHz, and saved on a PC computer running Clampex 10.4 (Molecular Devices). The extracellular solution used consisted of 133mM NaCl, 3mM KCl, 1mM MgCl2, 10mM HEPES, 2.5mM CaCl2, 10mM glucose, and 18.9 mM sucrose (pH 7.3 with NaOH, OSM=302). The patch pipettes (resistance of 3-10MΩ) were filled with an intracellular recording solution consisting of 133mM CsCl, 10mM HEPES, 5 mM EGTA, 1mM CaCl2, 1mM MgCl2, 4mM Mg-ATP, 0.4 mM Lithium-GTP, and 10mM Cs-gluconate with CsOH (pH 7.3 OSM=285).

Mechanical Activation of dissociated sensory neurons was performed by driving a heat polished blunt pipette tip (3-5mm) at an approximately 60 ° angle into the center of the cell using a micromanipulator mounted on a P841.20 piezoelectric translator (Physik Instrumente, Karlsruhe, Germany). A series of 10 increasing indentations (1μm increments) was applied to each cell. Cells that detached from the surface of the coverslip or lost the patch before the entire series of indentations were applied, were excluded. Cells were identified as responding only if the maximum evoked current exceeded 50pA. Fisher’s exact test with confidence level of 0.05 was used to test significance.

## Acknowledgements

We thank Miriam Lohnert, Annika von Seggern and Christina Steinmeyer-Stannek for excellent technical support (cell culture, *in situ* hybridization and southern blot techniques). We thank Thomas C. Südhof and Clifford J. Woolf for providing viral expression constructs. We thank Vladimir Benes from the Genomics Core Facility at EMBL for fruitful advice regarding deep sequencing experiments and the Nikon Imaging Center at the University of Heidelberg. This work was supported by the German Research Foundation (SCHR1523/2-1 to K.S.-S. and SFB-1158 to J.S.).

## Author contributions

J.S. and K.S.-S. conceived the study, designed experiments, and wrote the manuscript together with input and approval from all authors. K.S.-S. designed differentiation procedures and performed cell culture work, generated the TRKA wt and TRKA-PIEZO2 KO reporter lines, and conducted *in situ* hybridization, immunostainings and calcium imaging experiments. J.P. designed, conducted and analyzed the electrophysiological recordings, M.A. analyzed the deep sequencing data and performed calcium imaging experiments. C.R. carried out some cell culture work, *in situ* hybridizations and immunostainings and some calcium imaging experiments. S.L. picked and processed NOCL1 and NOCL2 cells for deep sequencing experiments. A.T.C., R.M.L. and M.S. designed and performed the mechanical stimulation experiments in the mouse wt and Piezo2 KO nociceptors.

## Data availability statement

All data will be made available upon publication.

## Expanded View Figure legends

**Figure EV1**

**Generation of hESC-derived nociceptor-like neurons**

A. Schematic drawing of the two lentiviruses used for infection of hESC-derived NCLCs. Lentivirus #1 expresses the transactivator rtTA driven by the ubiquitin-(Ub) promoter. Lentivirus #2 contains the tetracycline promoter site (tetO), were the transactivator can bind and induce –in the presence of doxycline (Dox)– the expression of *NEUROGENIN1* (*NGN1*), enhanced green fluorescence protein (EGFP) and the puromycin resistance cassette (Puro), all of which are linked via P2A and T2A sequences.

B. Live cell imaging showing the expression of EGFP in infected hNCLCs after 6 and 20 days in culture. Cells were exposed to doxycycline-containing medium for 10 days. Strong EGFP signal is observed after 6 days while the signal is almost absent after 20 days. Scale bar 20μm.

C. Time course of *NEUROGENIN1* (*NGN1*) and *NEUROGENIN2* (*NGN2*) transcript-levels after 0, 12, 24 and 31 days of NOCL3 differentiation. Data are shown as normalized counts ± SEM. N=3.

**Figure EV2**

**Generation of TRKA-tomato hESC reporter cell line**

A. Schematic illustration indicating the cloning strategy used to generate a hESC-*TRKA*-tomato reporter line. The tandem tomato sequence was introduced into the last exon (exon 17) of the *TRKA* locus before the stop codon (TAG) via a T2A linker sequence using CRISPR/Cas9 technology. Additionally, a blasticidin selection cassette was inserted in the *TRKA* 3’UTR area with the same cloning vector. To identify hESC clones with the correct genetic modification, a 5’ southern probe was designed recognizing a sequence around exon 16.

B. Southern blot done with DNA from 26 hESC clones, showing the presence of 4 positive clones indicated with an asterisk. Positive clones contain the wild type band at 2.6 kb as well as a band at 5.3 kb, indicating the correct incorporation of the reporter-sequence. The clone marked with an additional arrow was used for most differentiation procedures.

C. *In situ* hybridizations for tomato (red) and *TRKA* (green) expression in NOCL1 cells demonstrate co-expression of both markers in 98.7 ± 0.9% of cells. Nuclear marker DAPI in blue; scale bar 20 μm; N=3.

**Figure EV3**

**Calcium imaging experiments of various differentiated cells**

A. Representative fluorescence ratios (340nm/380nm) of hESC-derived low threshold mechanoreceptors (LTMRs, green) and NOCL3 cells (pink) loaded with the calcium indicator Fura2 after challenging them with mustard oil (200μM), menthol (500μM) or capsaicin (1μM). To visualize all neurons a stimulus of high K^+^ Ringer solution was added at the end. LTMRs did not respond to any stimulus besides high K^+^ Ringer solution. Data are shown as average ratio ± stdev (NOCL3 = 123 cells; LTMRs = 2 cells).

B. Representative pseudo-colour images of calcium responses in hESC-derived NCLCs differentiated after infection with a RUNX1 expressing lentivirus, before (baseline) and during capsaicin or mustard oil stimulation. High K^+^ Ringer solution was added at the end. Lower panel shows representative normalized fluorescence ratios (340nm/380nm) of the same cells, indicating absence of any capsaicin responders but a small fraction of mustard oil responders under these conditions.

C. Representative pseudo-colour images of calcium responses in hESC-derived NCLCs differentiated after infection with a *TRKA* expressing lentivirus, before (baseline) and during capsaicin or mustard oil stimulation. High K^+^ Ringer solution was added at the end. Lower panel shows representative normalized fluorescence ratios (340nm/380nm) of the same cells, indicating absence of any capsaicin responders and presence of a small fraction of mustard oil responders under these conditions.

**Figure EV4**

**Time-course of selected genes in NOCL3 neurons**

Heat map showing expression (normalized log2 counts) of gene transcripts during the course of NOCL3 development (samples were taken at 0, 12, 24 and 31 days, N = 3 per time point). Genes are categorized into functionally connected groups and those indicated bold have been analysed in more detail (neurotrans. = neurotransmission, transcript. = transcription).

**Figure EV5**

**Molecular characterization of NOCL1 and NOCL3 neurons**

A. Immunostaining and *in situ* hybridization of NOCL3 neurons with anti-ISL1 antibody (left and red) and a *TRKA*-probe (middle and green), showing that essentially all sensory neurons express *TRKA* (99.7 ± 0.3%; N=3).

B. *In situ* hybridization of NOCL1 neurons (upper panel) for *TRKA* (left and red) and *TRKB* (middle and green) and immunostaining plus *in situ* hybridization of NOCL3 neurons (lower panel) using an anti-ISL1 antibody (left and red) and a *TRKB* probe (middle and green) to show the presence of *TRKB*-transcripts in different NOCL cells. While NOCL1 neurons only rarely show presence of *TRKB*, the majority of NOCL3 neurons do express it.

C. Dual-color *In situ* hybridization of NOCL1 neurons (upper panel) for *TRKA* (left and red) and *TRKC* (middle and green) and immunostaining and *in situ* hybridization of NOCL3 neurons (lower panel) using an anti-ISL1 antibody (left and red) and a *TRKC* probe (middle and green) to show the presence of *TRKC* transcripts in different NOCL cells. The fraction of NOCL1 neurons expressing *TRKC* transcripts is minor, compared to NOCL3 neurons.

D. Comparison of MAFA transcript-levels between LTMRs and NOCL1–3 neurons, indicating that the NOCL3-differentiation procedure results in slightly higher MAFA transcript-levels compared to those of NOCL1 or NOCL2- differentiation procedures. LTMRs exhibit the highest MAFA levels (RNAseq MAFA transcript counts were normalized to *ISL1* transcript counts (N=3)).

E. Immunostaining of hESC-derived LTMRs and NOCL3 neurons using an anti-ISL1 antibody (left panel) and an anti-MAFA antibody (right panel), showing that the level of MAFA-protein in NOCL3 neurons is minor in comparison to LTMRs, despite the presence of some MAFA transcript in NOCL3 neurons (**d**) (arrow indicates a MAFA-positive NOCL3 cell). Scale bars in all Figs: 20μM.

**Figure EV6**

**Classification of hESC-derived sensory neuron subtypes**

Heat map showing the expression of transcripts from several genes classically used to categorize rodent sensory neurons into different categories (LTMR, NP = non-peptidergic, PEP = peptidergic), in hESC-derived LTMRs and NOCL cells. Gene counts were first normalized and then related to the pan-sensory marker *ISL1* and averaged. Depicted are center-scaled values (N=3).

**Figure EV7**

A. Analysis of action potential (AP) waveforms. The AP repolarization was analyzed as the deviation from a single exponential fit between the AP peak and the peak of the afterhyperpolarization potential as depicted in the inset and compared to the result obtained for derived LTMRs; LTMR (n=43), NOCL1 (n=109), NOCL2 (n=70), NOCL3 (n=90) and NOCL4 (n=87), *** (p≤0.0002), Mann-Whitney test.

B. Resting membrane potential V_m_ was measured in current clamp for LTMR (n=44), NOCL1 (n=108), NOCL2 (n=70), NOCL3 (n=90) and NOCL4 (n=71). n.s. (p≥0.1432), *** (p<0.0001), Mann-Whitney test.

C. Capacitance was measured as an estimation of cell size for LTMR (n= 42), NOCL1 (n=109), NOCL2 (n=70), NOCL3 (n=90) and NOCL4 (n= 99), *** (p≤ 0.0002), Mann-Whitney test.

D. Percentage of the ratio of the maximal TTX-resistant sodium current and the maximal sodium current, measured in current-voltage curves respectively. (A-D) In all graphs: each triangle in a scatter plot corresponds to one measured cell and mean ± SEM is depicted.

E. Normalized current-voltage curves (means ± SEM) for NOCL3 ≥ 24 days in culture (green, n=26) and ≥ 48 days in culture (dark green, n=29) in the presence of 300-500 nM TTX to isolate TTX-R currents. Inset: Area under the curve (AUC) ranging from −20 mV to 60 mV in 20 mV steps, **** (p<0.0001), red lines are the median.

F. Average (± SEM) current of neurons responding to a mechanical stimulation evoked by increasing displacement of the somatic cell membrane.

**Figure EV8**

**Molecular and electrophysiological characterization of PIEZO2 KO NOCL neurons**

A. Immunostaining and *in situ* hybridization of hESC-derived PIEZO2 knockout NOCL3 cells with an anti-ISL1 antibody (left and red) and a *TRKA*-probe (middle and green), indicating that almost all ISL1-positive cells are also *TRKA*- positive.

B. Immunostaining and *in situ* hybridization of hESC-derived PIEZO2 knockout NOCL3 cells with an anti-ISL1 antibody (left and red) and a *TRPV1*-probe (middle and green), indicating that almost all ISL1-positive cells are also *TRPV1*-positive. Scale bars 20 μm.

C. Somatic AP half width is similar between wild type and *PIEZO2* KO neurons, as assessed by Mann-Whitney test, * (p=0.0261), n.s. (p ≥ 0.0581). (D) Decay of the after hypolarization potential (AHP) τ_AHP_ is similar between wild type and *PIEZO2* KO neurons as assessed by Mann-Whitney test, n.s. (p ≥ 0.6226). In graphs **c**) and **d**), each marker in the scatter plot corresponds to one measured cell and mean ± SEM is depicted.

## References for materials and methods

Bajpai R, Chen DA, Rada-Iglesias A, Zhang J, Xiong Y, Helms J, Chang CP, Zhao Y, Swigut T, Wysocka J. CHD7 cooperates with PBAF to control multipotent neural crest formation. Nature 2010; 469:958–962.

Schrenk-Siemens K, Wende H, Prato V, Song K, Rostock C, Loewer A, Utikal J, Lewin GR, Lechner SG, Siemens J. PIEZO2 is required for mechanotransduction in human stem cell-derived touch receptors. Nat Neurosci. 2015; 1:10–16.

Wainger BJ, Buttermore ED, Oliveira JT, Mellin C, Lee S, Saber WA, Wang AJ, Ichida JK, Chiu IM, Barrett L, et al. Modelling pain in vitro using nociceptor neurons reprogrammed from fibroblasts. Nat Neurosci. 2015; 1:17–24.

Wende H, Lechner SG, Cheret C, Bourane S, Kolanczyk ME, Pattyn A, Reuter K, Munier FL, Carroll P, Lewin GR, et al. The transcription factor c-Maf controls touch receptor development and function. Science 2012; 335:1373–6.

Szczot M, Liljencrantz J, Ghitani N, Barik A, Lam R, Thompson JH, Bharucha-Goebel D, Saade D, Necaise A, Donkervoort S, et al. PIEZO2 mediates injury-induced tactile pain in mice and humans. Sci Transl Med 2018; 10.

Cavanaugh DJ, Chesler AT, Jackson AC, Sigal YM, Yamanaka H, Grant R, O’Donnell D, Nicoll RA, Shah NM, Julius D et al., Trpv1 reporter mice reveal highly restricted brain distribution and functional expression in arteriolar smooth muscle cells. J Neurosci. 2011; 13:5067–77.

Picelli S, Faridani OR, Bjorklund AK, Winberg G, Sagasser S, Sandberg R. Full-length RNA-seq from single cells using Smart-seq2. Nat. Protocols 2014; 9:171–181.

Chomczynski P, Sacchi N. Single-step method of RNA isolation by acid guanidinium thiocyanate-phorbol-chloroform extraction. Anal Biochem. 1987; 1:156–9.

Patro R, Duggal G, Love MI, Irizarry RA, Kingsford C. Salmon provides fast and bias-aware quantification of transcript expression. Nature Methods 2017; 14:417–419.

Andrews S. (2010). FastQC: a quality control tool for high throughput sequence data. Available online at: http://www.bioinformatics.babraham.ac.uk/projects/fastqc

Ewels P, Magnusson M, Lundin S, Käller M. MultiQC: summarize analysis results for multiple tools and samples in a single report. Bioinformatics 2016; 32:3047–3048.

Soneson C, Love MI, & Robinson MD, Differential analyses for RNA-seq: transcript-level estimates improve gene-level inferences. F1000Research 2015; 4:1521.

Love MI, Huber W, Anders S. Moderated estimation of fold change and dispersion for RNA-seq data with DESeq2. Genome Biology 2014; 15:550.

Pachitariu M, Stringer C, Dipoppa M, Schröder S, Rossi LF, Dalgleish H, Carandini M, Harris KD. Suite2p: beyond 10,000 neurons with standard two-photon microscopy. bioRxiv 2017; doi.org/10.1101/061507

Gu Z, Eils R, Schlesner M. Complex heatmaps reveal patterns and correlations in multidimensional genomic data. Bioinf. 2016; 18:2847–9.

## References

Bajpai, R., Chen, D.A., Rada-Iglesias, A., Zhang, J., Xiong, Y., Helms, J., Chang, C.P., Zhao, Y., Swigut, T., and Wysocka, J. (2010). CHD7 cooperates with PBAF to control multipotent neural crest formation. Nature 463, 958–962.

Basbaum, A.I., Bautista, D.M., Scherrer, G., and Julius, D. (2009). Cellular and molecular mechanisms of pain. Cell 139, 267–284.

Blanchard, J.W., Eade, K.T., Szucs, A., Lo Sardo, V., Tsunemoto, R.K., Williams, D., Sanna, P.P., and Baldwin, K.K. (2015). Selective conversion of fibroblasts into peripheral sensory neurons. Nat Neurosci 18, 25–35.

Bonnington, J.K., and McNaughton, P.A. (2003). Signalling pathways involved in the sensitisation of mouse nociceptive neurones by nerve growth factor. J Physiol 551, 433–446.

Borbiro, I., Badheka, D., and Rohacs, T. (2015). Activation of TRPV1 channels inhibits mechanosensitive Piezo channel activity by depleting membrane phosphoinositides. Sci Signal 8, ra15.

Caterina, M.J., Leffler, A., Malmberg, A.B., Martin, W.J., Trafton, J., Petersen-Zeitz, K.R., Koltzenburg, M., Basbaum, A.I., and Julius, D. (2000). Impaired nociception and pain sensation in mice lacking the capsaicin receptor. Science 288, 306–313.

Chambers, S.M., Qi, Y., Mica, Y., Lee, G., Zhang, X.J., Niu, L., Bilsland, J., Cao, L., Stevens, E., Whiting, P., et al. (2012). Combined small-molecule inhibition accelerates developmental timing and converts human pluripotent stem cells into nociceptors. Nat Biotechnol 30, 715–720.

Chesler, A.T., Szczot, M., Bharucha-Goebel, D., Ceko, M., Donkervoort, S., Laubacher, C., Hayes, L.H., Alter, K., Zampieri, C., Stanley, C., et al. (2016). The Role of PIEZO2 in Human Mechanosensation. N Engl J Med 375, 1355–1364.

Coste, B., Mathur, J., Schmidt, M., Earley, T.J., Ranade, S., Petrus, M.J., Dubin, A.E., and Patapoutian, A. (2010). Piezo1 and Piezo2 are essential components of distinct mechanically activated cation channels. Science 330, 55–60.

Davidson, S., Copits, B.A., Zhang, J., Page, G., Ghetti, A., and Gereau, R.W.t. (2014). Human sensory neurons: Membrane properties and sensitization by inflammatory mediators. Pain 155, 1861–1870.

Dirajlal, S., Pauers, L.E., and Stucky, C.L. (2003). Differential response properties of IB(4)-positive and -negative unmyelinated sensory neurons to protons and capsaicin. J Neurophysiol 89, 513–524.

Dubin, A.E., Schmidt, M., Mathur, J., Petrus, M.J., Xiao, B., Coste, B., and Patapoutian, A. (2012). Inflammatory signals enhance piezo2-mediated mechanosensitive currents. Cell Rep 2, 511–517.

Emery, E.C., Young, G.T., and McNaughton, P.A. (2012). HCN2 ion channels: an emerging role as the pacemakers of pain. Trends Pharmacol Sci 33, 456–463.

Ernsberger, U. (2009). Role of neurotrophin signalling in the differentiation of neurons from dorsal root ganglia and sympathetic ganglia. Cell Tissue Res 336, 349–384.

Fang, X., McMullan, S., Lawson, S.N., and Djouhri, L. (2005). Electrophysiological differences between nociceptive and non-nociceptive dorsal root ganglion neurones in the rat in vivo. J Physiol 565, 927–943.

Franck, M.C., Stenqvist, A., Li, L., Hao, J., Usoskin, D., Xu, X., Wiesenfeld-Hallin, Z., and Ernfors, P. (2011). Essential role of Ret for defining non-peptidergic nociceptor phenotypes and functions in the adult mouse. Eur J Neurosci 33, 1385–1400.

Gu, Y., and Gu, C. (2014). Physiological and pathological functions of mechanosensitive ion channels. Mol Neurobiol 50, 339–347.

Habib, A.M., Wood, J.N., and Cox, J.J. (2015). Sodium channels and pain. Handb Exp Pharmacol 227, 39–56.

Inserra, M.C., Israel, M.R., Caldwell, A., Castro, J., Deuis, J.R., Harrington, A.M., Keramidas, A., Garcia-Caraballo, S., Maddern, J., Erickson, A., et al. (2017). Multiple sodium channel isoforms mediate the pathological effects of Pacific ciguatoxin-1. Sci Rep 7, 42810.

Kalcheim, C., and Kumar, D. (2017). Cell fate decisions during neural crest ontogeny. Int J Dev Biol 61, 195–203.

Kim, S.E., Coste, B., Chadha, A., Cook, B., and Patapoutian, A. (2012). The role of Drosophila Piezo in mechanical nociception. Nature 483, 209–212.

Lallemend, F., and Ernfors, P. (2012). Molecular interactions underlying the specification of sensory neurons. Trends Neurosci 35, 373–381.

Le Pichon, C.E., and Chesler, A.T. (2014). The functional and anatomical dissection of somatosensory subpopulations using mouse genetics. Front Neuroanat 8, 21.

Lechner, S.G., Frenzel, H., Wang, R., and Lewin, G.R. (2009). Developmental waves of mechanosensitivity acquisition in sensory neuron subtypes during embryonic development. EMBO J 28, 1479–1491.

Li, C.L., Li, K.C., Wu, D., Chen, Y., Luo, H., Zhao, J.R., Wang, S.S., Sun, M.M., Lu, Y.J., Zhong, Y.Q., et al. (2016). Somatosensory neuron types identified by high-coverage single-cell RNA-sequencing and functional heterogeneity. Cell Res 26, 967.

Ma, Q., Fode, C., Guillemot, F., and Anderson, D.J. (1999). Neurogenin1 and neurogenin2 control two distinct waves of neurogenesis in developing dorsal root ganglia. Genes Dev 13, 1717–1728.

Momin, A., Cadiou, H., Mason, A., and McNaughton, P.A. (2008). Role of the hyperpolarization-activated current Ih in somatosensory neurons. J Physiol 586, 5911–5929.

Murthy, S.E., Loud, M.C., Daou, I., Marshall, K.L., Schwaller, F., Kuhnemund, J., Francisco, A.G., Keenan, W.T., Dubin, A.E., Lewin, G.R., and Patapoutian, A. (2018). The mechanosensitive ion channel Piezo2 mediates sensitivity to mechanical pain in mice. Sci Transl Med 10.

Pace, M.C., Passavanti, M.B., De Nardis, L., Bosco, F., Sansone, P., Pota, V., Barbarisi, M., Palagiano, A., Iannotti, F.A., Panza, E., and Aurilio, C. (2018). Nociceptor plasticity: A closer look. J Cell Physiol 233, 2824–2838.

Prato, V., Taberner, F.J., Hockley, J.R.F., Callejo, G., Arcourt, A., Tazir, B., Hammer, L., Schad, P., Heppenstall, P.A., Smith, E.S., and Lechner, S.G. (2017). Functional and Molecular Characterization of Mechanoinsensitive “Silent” Nociceptors. Cell Rep 21, 3102–3115.

Ranade, S.S., Woo, S.H., Dubin, A.E., Moshourab, R.A., Wetzel, C., Petrus, M., Mathur, J., Begay, V., Coste, B., Mainquist, J., et al. (2014). Piezo2 is the major transducer of mechanical forces for touch sensation in mice. Nature 516, 121–125.

Reichling, D.B., and Levine, J.D. (2009). Critical role of nociceptor plasticity in chronic pain. Trends Neurosci 32, 611–618.

Rose, R.D., Koerber, H.R., Sedivec, M.J., and Mendell, L.M. (1986). Somal action potential duration differs in identified primary afferents. Neurosci Lett 63, 259–264.

Rostock, C., Schrenk-Siemens, K., Pohle, J., and Siemens, J. (2018). Human vs. Mouse Nociceptors - Similarities and Differences. Neuroscience 387, 13–27.

Schrenk-Siemens, K., Wende, H., Prato, V., Song, K., Rostock, C., Loewer, A., Utikal, J., Lewin, G.R., Lechner, S.G., and Siemens, J. (2015). PIEZO2 is required for mechanotransduction in human stem cell-derived touch receptors. Nat Neurosci 18, 10–16.

Stantcheva, K.K., Iovino, L., Dhandapani, R., Martinez, C., Castaldi, L., Nocchi, L., Perlas, E., Portulano, C., Pesaresi, M., Shirlekar, K.S., et al. (2016). A subpopulation of itch-sensing neurons marked by Ret and somatostatin expression. EMBO Rep 17, 585–600.

Szczot, M., Liljencrantz, J., Ghitani, N., Barik, A., Lam, R., Thompson, J.H., Bharucha-Goebel, D., Saade, D., Necaise, A., Donkervoort, S., et al. (2018). PIEZO2 mediates injury-induced tactile pain in mice and humans. Sci Transl Med 10.

Szczot, M., Pogorzala, L.A., Solinski, H.J., Young, L., Yee, P., Le Pichon, C.E., Chesler, A.T., and Hoon, M.A. (2017). Cell-Type-Specific Splicing of Piezo2 Regulates Mechanotransduction. Cell Rep 21, 2760–2771.

Tan, C.H., and McNaughton, P.A. (2016). The TRPM2 ion channel is required for sensitivity to warmth. Nature 536, 460–463.

Tsantoulas, C., Lainez, S., Wong, S., Mehta, I., Vilar, B., and McNaughton, P.A. (2017). Hyperpolarization-activated cyclic nucleotide-gated 2 (HCN2) ion channels drive pain in mouse models of diabetic neuropathy. Sci Transl Med 9, eaam6072.

Usoskin, D., Furlan, A., Islam, S., Abdo, H., Lonnerberg, P., Lou, D., Hjerling-Leffler, J., Haeggstrom, J., Kharchenko, O., Kharchenko, P.V., et al. (2015). Unbiased classification of sensory neuron types by large-scale single-cell RNA sequencing. Nat Neurosci 18, 145–153.

Vandewauw, I., De Clercq, K., Mulier, M., Held, K., Pinto, S., Van Ranst, N., Segal, A., Voet, T., Vennekens, R., Zimmermann, K., et al. (2018). A TRP channel trio mediates acute noxious heat sensing. Nature 555, 662–666.

Viana, F., and Belmonte, C. (2008). Funny currents are becoming serious players in nociceptor’s sensitization. J Physiol 586, 5841–5842.

Viatchenko-Karpinski, V., and Gu, J.G. (2016). Mechanical sensitivity and electrophysiological properties of acutely dissociated dorsal root ganglion neurons of rats. Neurosci Lett 634, 70–75.

Wainger, B.J., Buttermore, E.D., Oliveira, J.T., Mellin, C., Lee, S., Saber, W.A., Wang, A.J., Ichida, J.K., Chiu, I.M., Barrett, L., et al. (2015). Modeling pain in vitro using nociceptor neurons reprogrammed from fibroblasts. Nat Neurosci 18, 17–24.

Wende, H., Lechner, S.G., Cheret, C., Bourane, S., Kolanczyk, M.E., Pattyn, A., Reuter, K., Munier, F.L., Carroll, P., Lewin, G.R., and Birchmeier, C. (2012). The transcription factor c-Maf controls touch receptor development and function. Science 335, 1373–1376.

Yarmolinsky, D.A., Peng, Y., Pogorzala, L.A., Rutlin, M., Hoon, M.A., and Zuker, C.S. (2016). Coding and Plasticity in the Mammalian Thermosensory System. Neuron 92, 1079–1092.

